# Toxicokinetics of a Pesticide Mixture in Earthworms Reveal Concentration-Dependent Bioaccumulation and Limited Interactions

**DOI:** 10.64898/2026.09.03.749139

**Authors:** Lisa Gollot, Juliette Faburé, Ghislaine Delarue, Laura Frattaroli, Sylvie Nélieu, Raphaël Royauté, Rémy Beaudouin

## Abstract

Despite their fundamental role in sustaining terrestrial biodiversity and ecosystem functioning, soils are increasingly contaminated by pesticides, often occurring as complex mixtures. Understanding chemical uptake, distribution, and elimination in soil organisms is essential for reliable risk assessment. We investigated the toxicokinetics of a binary mixture of imidacloprid and epoxiconazole in the earthworm *Aporrectodea caliginosa*. Single-substance experiments were conducted to estimate uptake and elimination rates, and allowed the implementation of single substance toxicokinetic models. A mixture experiment tested five concentration ratios to assess potential interactions. Epoxiconazole, an azole fungicide, exhibited rapid uptake and elimination (*k_u_* = 1.9 g_soil_⋅g_worm_^−1^⋅d^−1^; *k_e_* = 1.7 d^−1^), resulting in low bioaccumulation (BAF = 0.9), whereas imidacloprid, a neonicotinoid insecticide, accumulated slowly with low elimination (*k_u_* = 1.6 g_soil_⋅g_worm_^−1^⋅ d^−1^; *k_e_* = 0.12 d^−1^), producing high bioaccumulation (BAF = 20). The explicit inclusion of the 24-hour Petri dish depuration phase proved to be critical, especially for epoxiconazole, highlighting the influence of gut-clearance procedures on apparent kinetics. In accordance with only a small synergistic effect in the mixture, no toxicokinetic interactions were detected in the mixture; internal concentrations of each compound were consistent across ratios. Concentration-dependent patterns were observed, with indications of a saturation process at high exposure levels for imidacloprid, and an increased BAF at high concentrations for epoxiconazole. This finding questions the use of single, concentration-independent bioaccumulation factors for hazard assessment and highlights the value of integrated TKTD approaches.

## 1. Introduction

The widespread adoption of synthetic pesticides throughout the 20th and 21st centuries has fundamentally transformed agricultural practices, with global consumption reaching 2.2 million tonnes in 2019 (Bondareva & Fedorova, 2021). This chemical-intensive approach to crop protection has left a persistent environmental legacy, particularly evident in soil ecosystems where pesticide residues accumulate and persist. European monitoring data reveal that more than 80% of agricultural soils contain detectable pesticide residues (Silva et al., 2019), while comprehensive French surveys indicate universal contamination across all tested agricultural sites (Pelosi et al., 2021). The scale of this contamination, with approximately 40,000 tonnes of pesticides applied annually in France alone (MTE, 2022), means that soil-dwelling organisms are routinely exposed to complex chemical mixtures rather than single compounds.

As soil engineers, earthworms represent key organisms for soil functioning and constitute the most important animal biomass in most terrestrial ecosystems (Lavelle et al., 2006). Their ecological niche makes them particularly vulnerable to pesticide contamination, as soil serves as both their habitat and their food source. Earthworms are therefore exposed to pesticides through multiple pathways: dermal contact during burrowing, ingestion of contaminated soil particles and organic matter, and uptake from pore water (Jager et al., 2003). The toxicokinetics of pesticides in earthworms, encompassing absorption, distribution, metabolism, and excretion (ADME) processes, determines internal exposure levels and ultimately influences toxicodynamic (TD) responses.

Toxicokinetic interactions in pesticide mixtures can occur through multiple pathways and at different temporal phases of exposure. During uptake phases, compounds may compete for the same transport mechanisms, modify membrane permeability, or saturate binding sites, leading to altered absorption rates. During elimination phases, metabolic interactions become particularly important, as compounds can competitively inhibit or induce biotransformation enzymes, modify tissue distribution, or interfere with efflux processes (Cedergreen, 2014). These interactions can result in either enhanced or reduced bioaccumulation depending on the specific mechanisms involved, with consequences that vary over different exposure durations.

Despite growing recognition of mixture toxicity, most ecotoxicological studies continue to focus on single compounds, with limited understanding of how co-exposure affects pesticide toxicokinetics. This knowledge gap is particularly concerning given that agricultural soils typically contain multiple pesticide residues simultaneously (Silva et al., 2019). Moreover, exposure is inherently dynamic: external concentrations fluctuate over time due to degradation, sorption, and organism-driven processes, while internal concentrations result from the continuous interplay between uptake and elimination. As a consequence, snapshot measurements of internal concentrations provide only a partial view of exposure, making it difficult to disentangle the mechanisms driving mixture effects.

In this context, modeling approaches provide a powerful framework to integrate these dynamic processes and quantitatively describe the links between external exposure and internal concentrations over time. Toxicokinetic (TK) modeling in earthworms typically employs one-compartment models that describe chemical bioaccumulation through uptake and elimination rate constants (*k_u_* and *k_e_*), providing bioaccumulation factors (BAF) and internal concentration dynamics over time (Jager et al., 2003). These models assume first-order kinetics and allow quantification of steady-state concentrations and half-lives, forming the foundation for understanding chemical fate in earthworm tissues. However, when multiple pesticides co-occur, these fundamental TK parameters can be significantly altered through various interaction mechanisms, potentially invalidating single-compound predictions.

Epoxiconazole (EPX) and imidacloprid (IMD), represent priority contaminants identified in French agricultural soils (Pelosi et al., 2021). Epoxiconazole, a lipophilic triazole fungicide (log*K_ow_* = 3.3, Solubility = 7.1 mg/L at 20°C at pH 7) that inhibits sterol biosynthesis, and imidacloprid, a neonicotinoid insecticide targeting nicotinic acetylcholine receptors (Chen et al., 2021; Lewis et al., 2006), exemplify the fungicide-insecticide combinations commonly found in agricultural environments. Despite their EU ban, environmental persistence ensures continued exposure risks for soil organisms.

Both pesticides demonstrate significant toxicity in earthworms. Epoxiconazole exposure causes substantial reductions in growth and reproductive output (EC_50, reproduction_ = 126.8 mg/kg ; NOEC_Juvenile growth_ = 9.3 mg/kg) (Bart et al., 2019; Gollot et al., 2026b; Pelosi et al., 2016), while triggering oxidative stress and energy depletion indicative of metabolic costs (Givaudan et al., 2014; Pelosi et al., 2016). Imidacloprid similarly impairs earthworm reproduction (EC_50_ = 0.55 mg/kg) and juvenile growth (NOEC = 0.28 mg/kg) (Gollot et al., 2026b), with concentration-dependent effects on activity and cast production (Dittbrenner et al., 2010, 2011; Van Loon et al., 2022; Wang et al., 2019).

The theoretical basis for imidacloprid-epoxiconazole toxicokinetic interactions centers on metabolic interference mechanisms. Triazole fungicides such as epoxiconazole are well-documented inhibitors of cytochrome P450 monooxygenases, key enzymes involved in xenobiotic biotransformation (Bass et al., 2015; Cedergreen et al., 2006; Puinean et al., 2010). By inhibiting these enzymes, triazoles can alter the elimination kinetics of co-occurring compounds, potentially decreasing elimination rate constants and increasing bioaccumulation factors. In *Daphnia magna*, such metabolic inhibition enhanced esfenvalerate toxicity by up to fourfold (Cedergreen et al., 2006) and increased *α*-cypermethrin toxicity threefold (Gottardi et al., 2017), suggesting that similar magnitude changes in internal concentrations might occur in earthworms.

While moderate hydrophilicity of imidacloprid (log*K_ow_* = 0.57, Solubility = 610 mg/L at 20°C at pH 7) might suggest that it can be excreted with limited metabolic transformation, evidence from insect species indicates partial metabolism via cytochrome P450 enzymes (Chen et al., 2018; Chen et al., 2019). Cytochrome P450 overexpression has been linked to imidacloprid resistance in multiple insect species (Elzaki et al., 2017; Kaplanoglu et al., 2017; Liang et al., 2015). Comparable metabolic pathways likely exist in earthworms, as suggested by studies reporting cytochrome P450 activity and induction, but their role in pesticide biotransformation remains less well characterized than in insects (Sanchez-Hernandez et al., 2014).

To address this knowledge gap, this study aims to characterize the toxicokinetics of epoxiconazole and imidacloprid in earthworms exposed to each compound individually and in mixture. Using bioaccumulation experiments coupled with toxicokinetic modeling, we examine uptake rates, elimination kinetics, and accumulation patterns to elucidate potential toxicokinetic interactions that may underlie mixture effects in this environmentally relevant species. This mechanistic approach will provide insights into the biological processes governing pesticide mixture toxicity and contribute to more robust environmental risk assessment frameworks.

To this overall goal, first, we investigate the individual toxicokinetic behaviors of epoxiconazole and imidacloprid in earthworms and examine how their contrasting physicochemical properties influence bioaccumulation patterns. We hypothesize that epoxiconazole and imidacloprid will exhibit distinct toxicokinetic profiles reflecting their contrasting physicochemical properties, with epoxiconazole showing higher bioaccumulation potential compared to imidacloprid. Second, we determine whether co-exposure to epoxiconazole alters imidacloprid bioaccumulation kinetics through metabolic interference, potentially explaining mixture toxicity mechanisms. We predict that co-exposure will result in an increased internal concentration of imidacloprid compared to single-substance exposure, potentially due to reduced biotransformation and/or elimination.

## 2. Material & Methods

### 2.1 Earthworm, soil and pesticides

Individuals used for this study were bred in the laboratory from individuals initially collected in late 2023 and early 2024 from a natural population in a permanent grassland in Versailles (48°48′ N, 2°5′ E) where no pesticides have been applied for more than 25 years. Individuals were identified using a binocular microscope and an identification key. Individuals are bred at 18°C in 1 L vessels in groups of five individuals using a soil sampled from this same meadow, a loamy soil texture (based on the texture definition of the Food and Agriculture Organization of the United Nations, FAO), to be as close as possible to the conditions of the natural population. More information on the soil is available in the Supplementary information Table S1.1. The soil was collected from the top 0-20 cm, air-dried and crushed to pass a 2 mm mesh. Soil culture is kept with a soil water-holding capacity (WHC) of 60-70% and individuals are fed with horse dung frost and defrosted twice before being milled (> 0.5 mm) as presented in Lowe & Butt (2005). The amount of 6 g of dry horse dung per group of five are given with a WHC of 60-80% every month. As recommended by Bart et al. (2018), ten individuals from the natural population were barcoded, showing individuals come from two lineages of the *A. caliginosa* species (see Supplementary information Figure S1.1).

For all experiments, the culture soil was used and contaminations were performed using imidacloprid and epoxiconazole sold as analytical standard by Sigma-Aldrich (purity ≥ 98.0% and 98.75% for IMD and EPX, respectively). Due to the very low water solubility of epoxiconazole (7.1 mg/L) and the relatively high tested concentrations, it was not possible to spike the soil with a water solution containing both epoxiconazole and imidacloprid. The alternative use of a co-solvent was avoided, as they are often toxic to earthworms, and the required quantity would have been excessive. One day prior to the start of the tests, the soil contamination was carried out in two stages: first, a dry mixture of epoxiconazole and soil at 20% of its WHC was prepared, followed by spiking the soil with a water solution of imidacloprid. Soil samples were collected at the start of each experiment for every soil batch prepared.

### 2.2 Toxicokinetic experiments

#### 2.2.1 Single substance exposure

A single substance toxicokinetic experiment was conducted following an adapted OECD Test Guideline 317 protocol (OECD, 2010) with destructive sampling to determine uptake and elimination rates of each pesticide in earthworms. The experimental design consisted of two sequential phases: a 21-day uptake phase where earthworms were exposed to pesticide-spiked soil, followed by a 21-day elimination phase where individuals were transferred to control soil for depuration. Sampling was conducted at eight timepoints during each phase: days 1, 2, 3, 7, 10, 14, 17, and 21 for uptake, and days 22, 23, 24, 28, 31, 35, 38, and 42 for elimination.

Each replicate consisted of a single adult earthworm maintained in an individual plastic vessel (11 × 7.7 × 4.5 cm) containing 200 g dry weight of soil adjusted to 70% water holding capacity. This design enabled individual-level measurements of both growth and internal concentrations throughout the experiment. Organisms were fed 3 g dry weight of horse dung per individual every 14 days and maintained at 18°C in a temperature-controlled environment. Individual body weights were recorded at exposure initiation and at respective sampling timepoints. At each timepoint, four individual earthworms were starved for approximately 24 hours on wet filter paper in Petri dishes to clear gut contents, then weighted and sacrificed to be stored at -80°C for subsequent chemical analysis. When the 24 hours were not enough to clear gut content, individuals were massaged to clear the remaining of the gut content.

Test concentrations in soil were set at 100 ng/g for imidacloprid and 1000 ng/g for epoxiconazole, selected to avoid acute toxicity while ensuring analytical quantifiability. These concentrations represent environmentally relevant exposure levels, approximating maximum concentrations reported in French agricultural soils (Pelosi et al., 2021), providing an optimal balance between analytical detectability and ecological relevance.

Given the rapid uptake and elimination already observed for epoxiconazole in earthworms (Bart et al., 2020), additional early sampling points were included to improve the estimation of kinetic parameters and reduce potential bias associated with the depuration phase. For these timepoints, earthworms were directly massaged to remove gut content without a depuration period in Petri dishes prior to storage at -80°C. Three additional sampling points were included per pesticide (0.25, 1, and 3 days for epoxiconazole; 3, 7, and 14 days for imidacloprid), each with three replicates.

#### 2.2.2 Exposure to the mixture

Adult earthworms were exposed for 28 days to a range of mixtures of imidacloprid and epoxiconazole varying both in concentration levels and in component ratios. The 28-day exposure period in the mixture experiment was inherited from the original reproduction-oriented design (Gollot et al., 2026b). In the context of the present study, it also offered the advantage of providing validation points for the single-substance TK models beyond the 21-day duration of the OECD-based experiment. Earthworms were maintained in pairs within experimental cosms containing 200 g of soil and were fed with 3 g of food per individual every 14 days. Body weights were recorded at the start of the exposure, after 14 days, and at the end of the 28-day period. Following exposure, earthworms were transferred to Petri dishes lined with moist filter paper and kept fasting for 26 hours to allow gut clearance. After this depuration phase, individuals were weighed again, sacrificed, and stored at -80 °C for chemical analysis. As for the first experiments, when 24 h was not sufficient to completely clear the gut, individuals were gently massaged to remove the remaining gut contents. Two effect isoboles were tested with five ratios of concentration each: 1:0 (EPX only), 3:1, 1:1, 1:3, 0:1 (IMD only). The first isobole corresponds to concentrations relatively comparable to the exposure concentrations in the single substance experiments while the second corresponds to much higher concentrations (x18).

### 2.3 Chemical analysis

The chemical analysis method is fully described in the Supporting Information Section S2.5. Briefly, aliquots of 5 g soil or 300 mg earthworms were spiked with deuterated internal standards and extracted by a QuEChERS method, using sodium acetate and magnesium sulphate. A further purification step was performed by freezing the earthworm extract to allow protein precipitation, and for both matrices using d-SPE sorbents containing C18 and PSA phases.

The samples were then analysed by on-line Solid Phase Extraction Ultra-High Performance Liquid Chromatography coupled via an ElectroSpray Interface to a triple quadrupole Mass Spectrometer (SPE-UHPLC-ESI-MS/MS). After injecting 600 µL on the SPE cartridge, the chromatographic separation was performed using a water/acetonitrile (both acidified) gradient on a C18 column. The detection was performed in positive mode using specific transitions for EPX and IMD by multiple reaction monitoring.

### 2.4 Toxicokinetic model

Two toxicokinetic model structures were tested to describe the dynamics of internal pesticide concentrations in earthworms: a one-compartment and a two-compartment model. The two-compartment structure was considered to account for potential differential distribution of compounds within the organism, particularly for epoxiconazole, a lipophilic substance that may partition into lipid-rich tissues. For consistency and comparability between compounds, the same two-compartment structure was also applied to imidacloprid. The one- and two-compartment models are hereafter referred to as “C1” and “C2”, respectively. Both model structures explicitly account for growth dilution.

In addition, two exposure scenarios were tested to account for the depuration phase prior to chemical analysis. Before freezing, earthworms were transferred to Petri dishes lined with moist filter paper for 24 hours to allow gut clearance and avoid measuring soil residues in the digestive tract. During this period, worms were no longer in direct contact with contaminated soil; however, residual exposure could still occur through ingestion of contaminated soil particles retained in the gut. To evaluate the influence of this potential pathway, two alternative exposure scenarios were implemented, differing in whether uptake was assumed to continue during the depuration phase.

In each model, the internal contaminant concentration (*C*_int_) is then modeled by a differential equation integrating three processes: (i) uptake from soil, which is proportional to *C*_ext_ via the uptake rate constant (*k_u_*); (ii) elimination from the organism, which follows first-order kinetics with rate constant (*k_e_*); and (iii) growth dilution, expressed as a term proportional to the growth rate and the current internal concentration. This last process reduces the concentration, reflecting the increase in biomass. All state variables and parameters are listed Table 1.

**Table 1:**
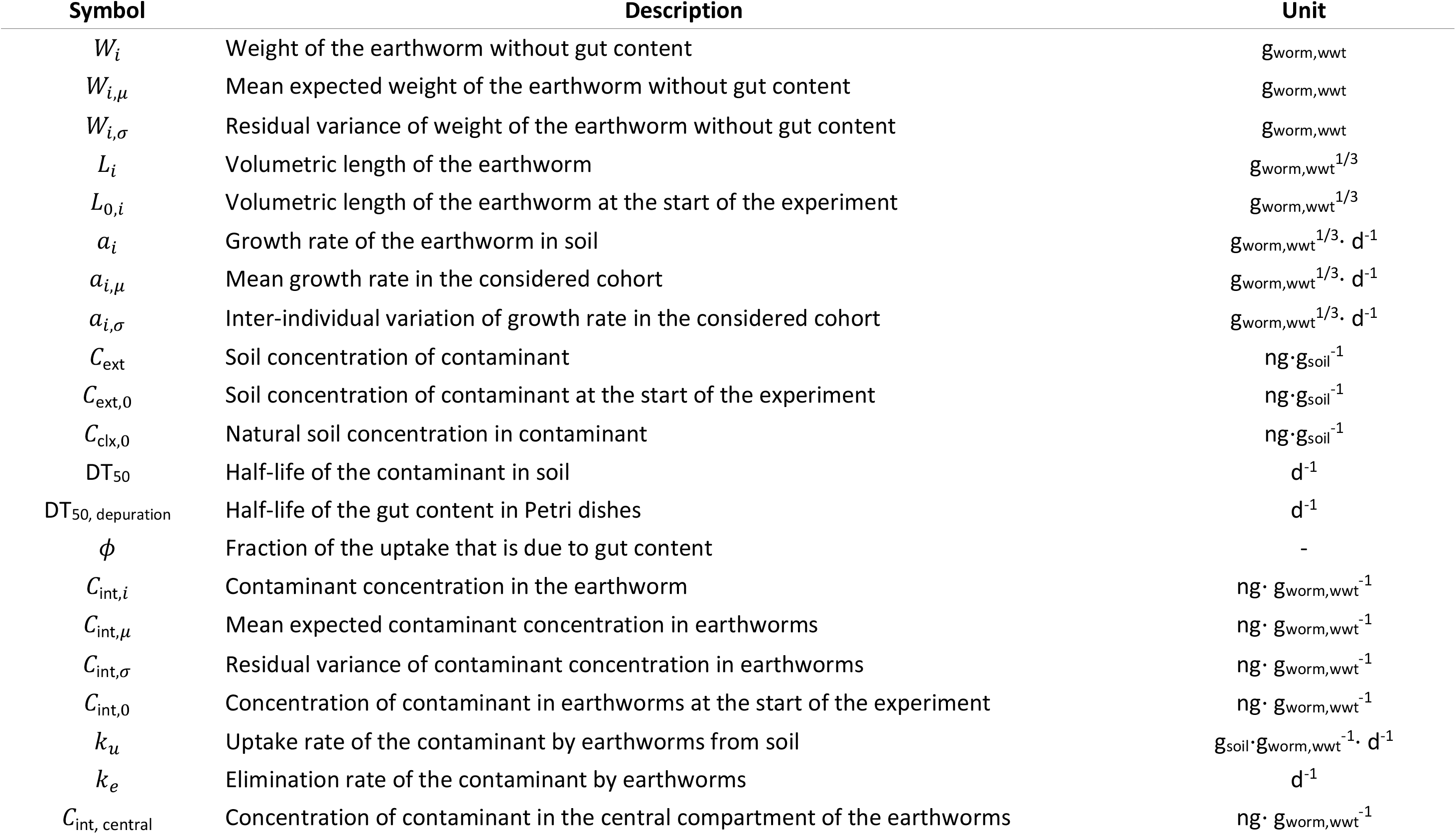
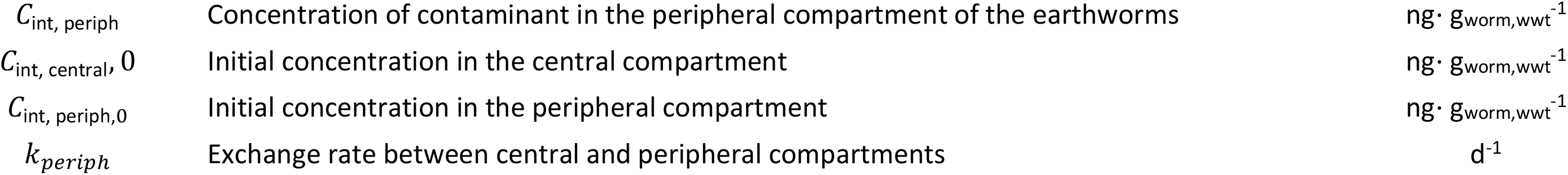
State variables and parameters list.

In the one-compartment model (C1), the internal concentration (*C*_int_) changes over time as a balance between uptake from soil (*C*_ext_), elimination to the external medium, and dilution by growth. Because individual body weights were available throughout the experiment, a hierarchical model for growth was implemented on the same principle as in Gollot et al. (2025). This dynamic is governed by Equation 1.

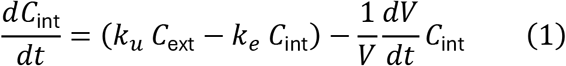

where *k_u_* and *k_e_* are the uptake and elimination rates of the contaminant by the earthworm, *L* is the structural length of the earthworm, and *a_i_* the individual growth rate.

In the two-compartment model (C2) model, the earthworm body is considered as consisting of a central compartment (e.g., blood or well-perfused tissues) and a peripheral compartment (e.g., storage or poorly perfused tissues), with bidirectional exchanges between them (*k*_periph_). Uptake from soil occurs only into the central compartment, from which the substance can either be eliminated to the external medium or distributed to the peripheral compartment. The peripheral compartment acts as a storage site that can delay elimination and buffer fluctuations in exposure, which is particularly relevant for hydrophobic compounds expected to partition into lipid-rich tissues. Such a structure allows us to account for a biphasic accumulation and elimination pattern, typically characterized by a fast initial phase driven by the central compartment and a slower phase reflecting the exchange with the peripheral compartment. The internal concentration is now described by Equation 2, where the dynamics of the central and peripheral compartments are jointly expressed, enabling a more mechanistic representation of toxicokinetics for lipophilic pesticides in earthworms.

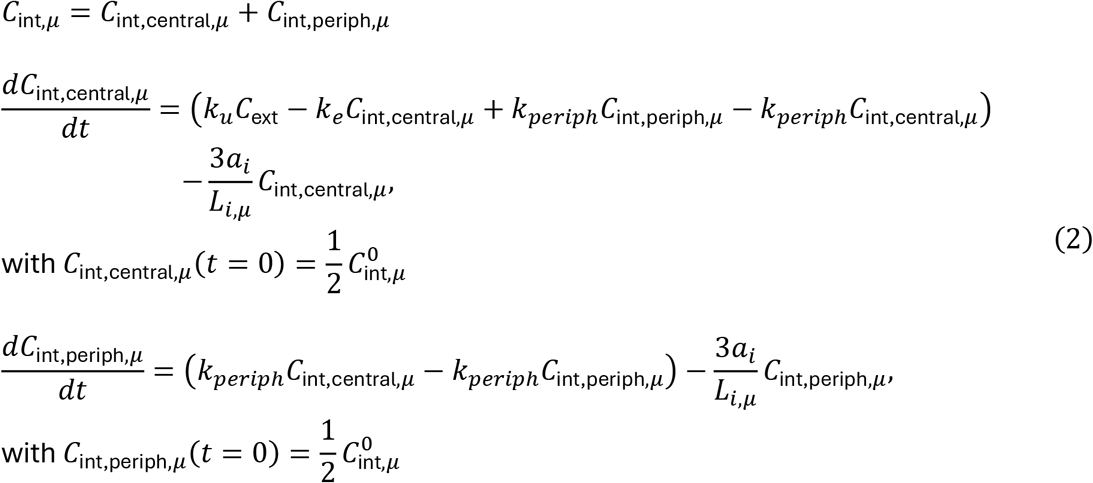

The external contaminant concentration in soil (*C*_ext_) is described by first-order dissipation, parameterized by the pesticide-specific half-life (DT_50_) (Equation 3). For epoxiconazole and imidacloprid, the DT_50_ values are 353.3 and 187 days, respectively. The initial concentration corresponds to the spiked soil, which decreases over time following first-order kinetics, and after 21 days is set equal to the residual concentration measured in natural soil. For the phase when earthworm are put on wet filter paper in Petri dishes for depuration, we consider that no exposure occurs and therefore *C*_ext_ is set to zero.

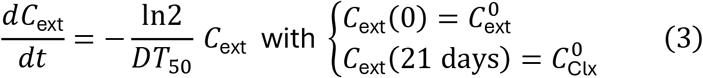

To account for potential residual exposure during the depuration phase, we tested two alternative exposure scenarios : (i) no exposure occurring in Petri dishes and (ii) assuming a digestive contribution of % of the total uptake when earthworms are in Petri dishes with contaminated soil still in their gut content (models noted with a “P”). Depuration of the earthworm gut content was modeled as a first-order decay, informed by a small supporting experiment: ten earthworms were kept individually in 200 g of Closeaux soil with 3 g of horse dung for one week, then transferred to Petri dishes lined with moist filter paper and weighed at regular intervals (model details in Supplementary information Section S2.3). After 32 hours, the worms were gently massaged to void the remaining gut contents and weighed again, providing data to estimate the gut-clearance rate.

To evaluate the influence of the depuration phase on toxicokinetic parameter estimation, different approaches were implemented depending on the compound. For imidacloprid, a reduced two-compartment model (C2N) was developed by assuming no net exchange during the depuration phase (uptake and elimination rates set to zero), and its performance was compared to the C2 model using the Bayesian Information Criterion (BIC). For epoxiconazole, the impact of the depuration phase was assessed by comparing the bioaccumulation factor estimated from the TK model with the steady-state BAF (BAF_ss_) calculated directly from measured concentrations after depuration, following OECD TG 317. This approach allowed us to quantify the bias introduced by neglecting the depuration phase for compounds with rapid toxicokinetics.

A linear model fitted to paired measurements of earthworm body weight with and without gut content indicated that gut content represents approximately 15% of fresh weight (*W*_without gut content_ = 0.85 × *W*_with gut content_, *R*^2^= 0.992), and was subsequently used to convert measurements with gut content into gut-free body weight for consistency across datasets (details are provided in Supplementary information Section 2.2).

Then, the structural length of the earthworm (*L* = *V*^1/3^) is related to the weight without gut content (*W*) through an allometric relationship with a 1/3 power scaling. Both internal concentration and individual weight are assumed to follow probability distributions, with internal concentration following a log-normal distribution of a mean *C*_int,*μ*_, and a variance *C*_int,*σ*_, and weight normally distributed around a mean, *W_μ_*, with a specified standard deviation, *W_σ_* (Equation 4).

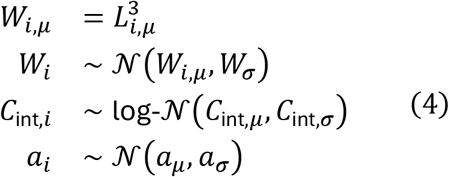

Growth in soil is described by a growth rate parameter drawn from a normal distribution (*a_i_*), which is set to zero during fasting phases in Petri dishes, where no growth is assumed (Equation 5) (Gollot et al., 2025).

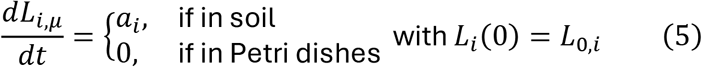

Model calibration was performed with a bayesian inference software, MCSim, using 3 chains and 100,000 iterations. Parameter values were inferred by Bayesian numerical (Bois, 2009). Other statistical analyses were performed using the software R.

Large and weakly informative priors were assigned to the toxicokinetic rate constants in order to cover a wide range of plausible values. Specifically, the uptake (*k_u_*), elimination (*k_e_*), and inter-compartmental exchange rates (*k_perip_*_ℎ_) were given large log-uniform priors, ensuring that the posterior estimates would be primarily informed by the data rather than overly restrictive assumptions. For the growth process, truncated normal priors were specified for both the mean growth rate and its inter-individual variation, reflecting prior biological knowledge of adult earthworm growth while avoiding unrealistic values. Priors used for each model are given in Supplementary information Section S3.1.

Measurement processes were also integrated in the model: observed body weights were assigned truncated normal distributions to ensure positivity and consistency with biological constraints, while internal concentrations were assumed to follow truncated log-normal distributions, which guarantee non-negative values.

Posterior distributions were drawn using the last 10,000 iterations of each chain. BAF_k_ were calculated as the ratio between *k_u_* and *k_e_*, using the last 10,000 iterations from each MCMC chain. Model comparison was based on the Bayesian Information Criterion (BIC), which balances goodness-of-fit and model complexity by penalizing models with a higher number of parameters.

### 2.5 Toxicokinetic simulations for mixture exposures

Simulations were conducted using the toxicokinetic models previously calibrated on single-compound exposure experiments to predict internal concentrations of both pesticides in earthworms exposed to mixtures. These simulations assume no interaction between compounds and therefore provide a null expectation based on independent toxicokinetics. Predicted concentrations were compared to experimental data obtained for both single-compound exposures and mixture treatments. In particular, comparisons were performed across the different mixture ratios tested to assess whether co-exposure to epoxiconazole modified the toxicokinetics of imidacloprid. Deviations between model predictions and observed concentrations would be interpreted as evidence of potential toxicokinetic interactions.

To account for dilution due to growth, the individual growth rate of each earthworm during the mixture exposure was estimated by linear regression and directly implemented in the model.

## 3 Results

### 3.1 Depuration dynamic

Gut clearance in earthworms was assumed to follow first-order kinetics, with an estimated DT_50,depuration_ of 4.58 hours (CI [3.6 ; 6.2]) (Figure 1). The gut content represented 16.7% (CI [15.4; 18]) of the total wet weight prior to depuration. This estimate is consistent with the previously established relationship based on a larger dataset comparing earthworms with and without gut content, supporting the robustness of this value. Two earthworms were excluded from the analysis as they exhibited an increase in weight during the experiment, likely due to water ingestion, and therefore did not undergo normal depuration.

**Figure 1:**
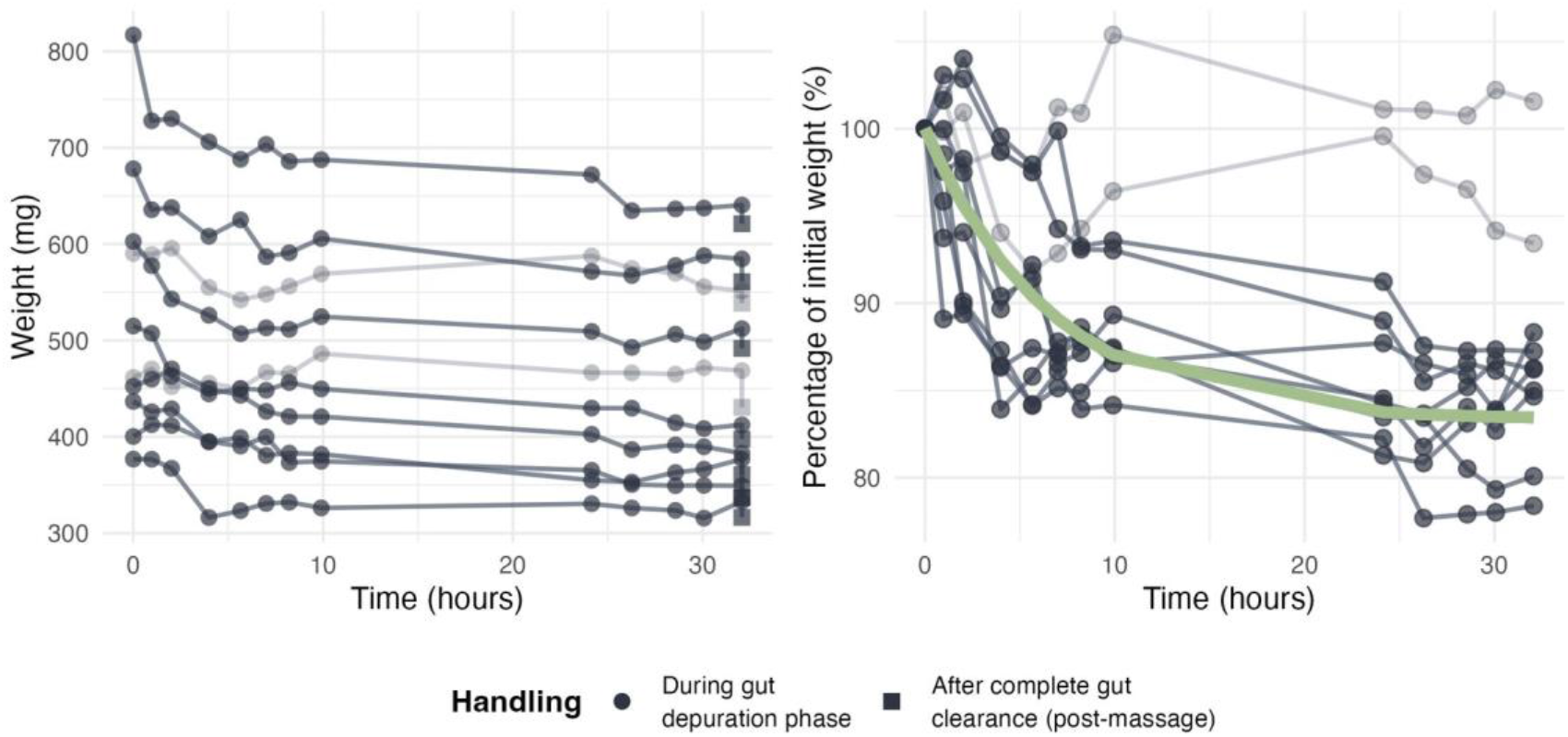
Evolution of earthworm weight (left) and the corresponding percentage of initial weight (right) throughout time in Petri dishes. The green line corresponds to the prediction of the model. Weight after complete gut clearance (post massage) are described with squared points. Transparent individuals were excluded from the analysis.

### 3.2 Single exposures

The single-substance toxicokinetic experiments were successfully conducted, with no mortality observed throughout the uptake and elimination period. Earthworms weight used in the first single substance experiments had a median of 619.9 mg (Q1 = 522.3 mg; Q3 = 701.3 mg; *n* = 73) for IMD, and 596.2 mg (Q1 = 514.0; Q3 = 668.2; *n* = 71) for EPX. A total of three individuals out of 146 were excluded as outliers prior to data analysis: two earthworms at the 35-day time point for epoxiconazole and one earthworm at the 17-day time point for imidacloprid. Their internal concentrations were markedly inconsistent with both model predictions and co-sampled replicates, deviating by approximately 100-fold and 8-fold, respectively, suggesting potential analytical or sampling artefacts. For epoxiconazole, marked differences in measured internal concentrations were observed between earthworms subjected to depuration in Petri dishes and those for which gut contents were removed by direct massage at comparable sampling time points. This discrepancy indicates that significant elimination occurred during the depuration phase, leading to lower apparent internal concentrations in depurated individuals. In contrast, such differences were less pronounced for imidacloprid, consistent with its slower toxicokinetic behavior and lower sensitivity to elimination during the depuration phase.

All tested models fitted correctly to the data, and the growth module enabled 100% of body weight predictions to fall within the 2-fold change region (see Supplementary information Section S3).

In the case of imidacloprid, the two-compartment model (C2) provided the best fit to the data, with the lowest BIC value (Table 2). Overall, 100% of predictions fell within the 5-fold change region and 90.3% within the 2-fold change region (Figure 2), confirming the excellent predictive performance of the model, with individual trajectories closely matching observed data points (Figure 4). Across the two models tested, however, the initial phase of uptake was consistently slightly underestimated and the second phase of elimination was slightly overestimated (see Supplementary information Figure S3.11). Additionally, exposure through the gut route in Petri dishes appears unlikely, as reflected by the higher BIC value for the C2P model. The C1P model was also tested, but the parameter, and consequently all remaining parameters, could not be reliably estimated (see Supplementary information Section S3.6). The estimated toxicokinetic parameters of all tested models are summarized in Table 3.

**Figure 2:**
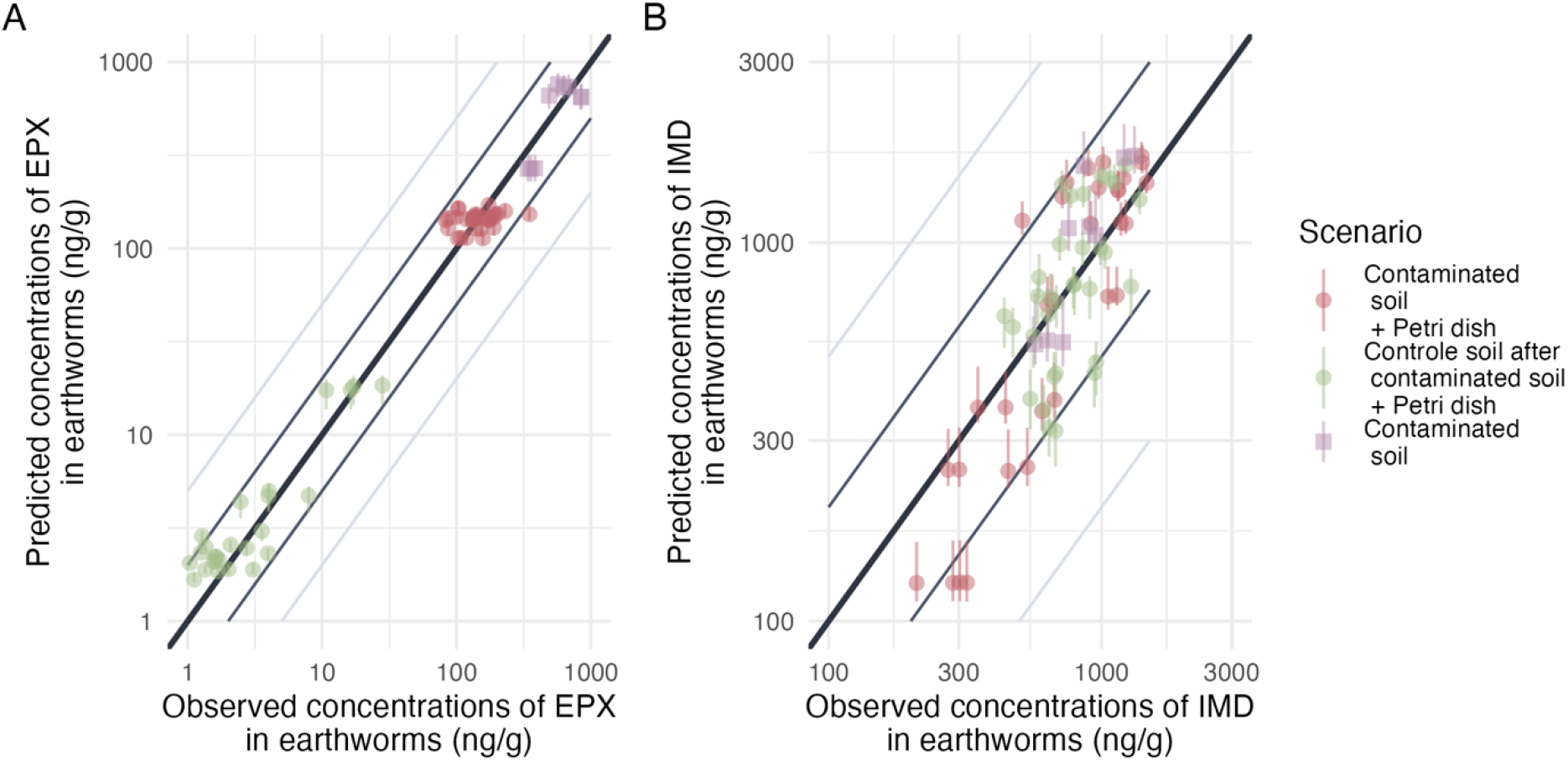
Predicted versus observed values of (A) imidacloprid (one-compartment model) and (B) epoxiconazole (two-compartment model, exposure through the gut route in Petri dishes) internal concentrations. Points are colored by exposure scenario: red indicates earthworms exposed only to contaminated soil (uptake phase), green indicates earthworms transferred from contaminated to clean soil (elimination phase) and violet indicates earthworms which were massaged instead of put in Petri dishes for 24h for depuration of their gut content. Grey and light-grey lines represent the 2-folds and 5-fold changes, respectively. The bold black line represents the identity line.

**Figure 3:**
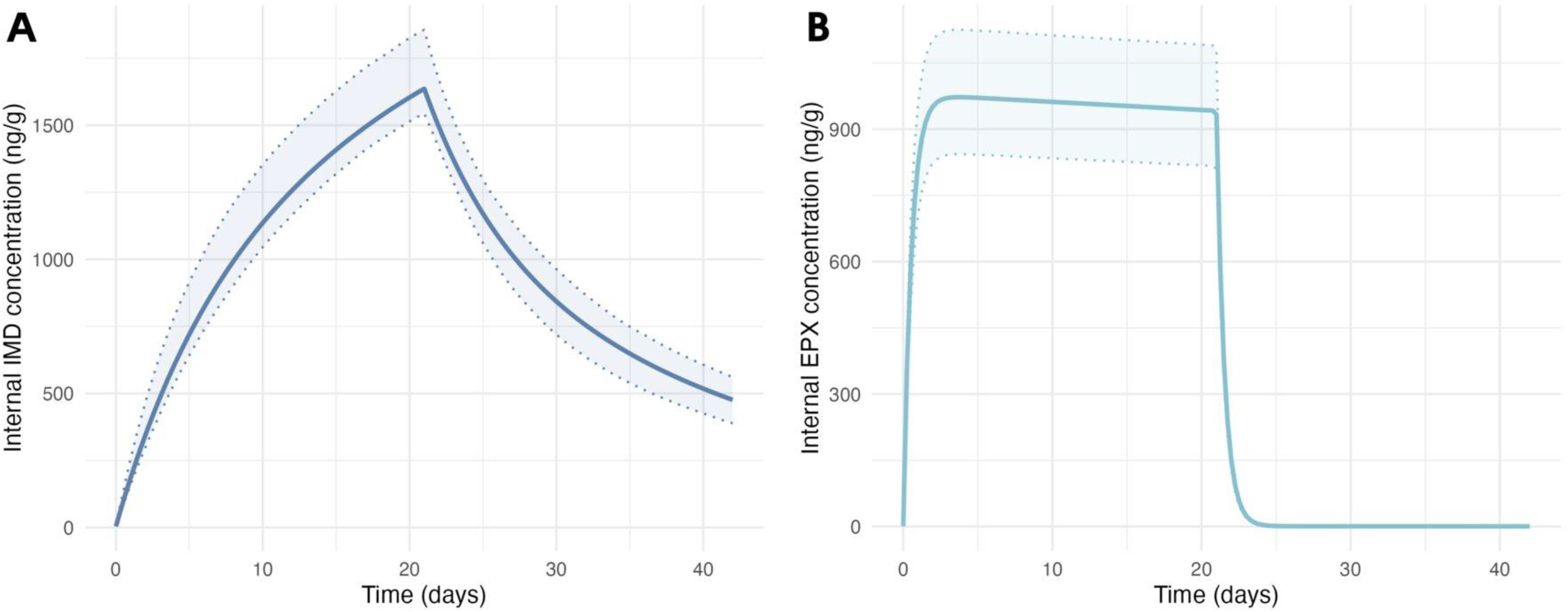
Estimated toxicokinetic of (A) imidacloprid (IMD) and (B) epoxiconazole (EPX) concentration in earthworms exposed to 100 ng/g of IMD and 1000 ng/g of EPX, respectively. (IMD in dark blue and EPX in light blue). The curve represents the estimated toxicokinetic curve and the transparent band corresponds to the credibility interval of the model.

**Figure 4:**
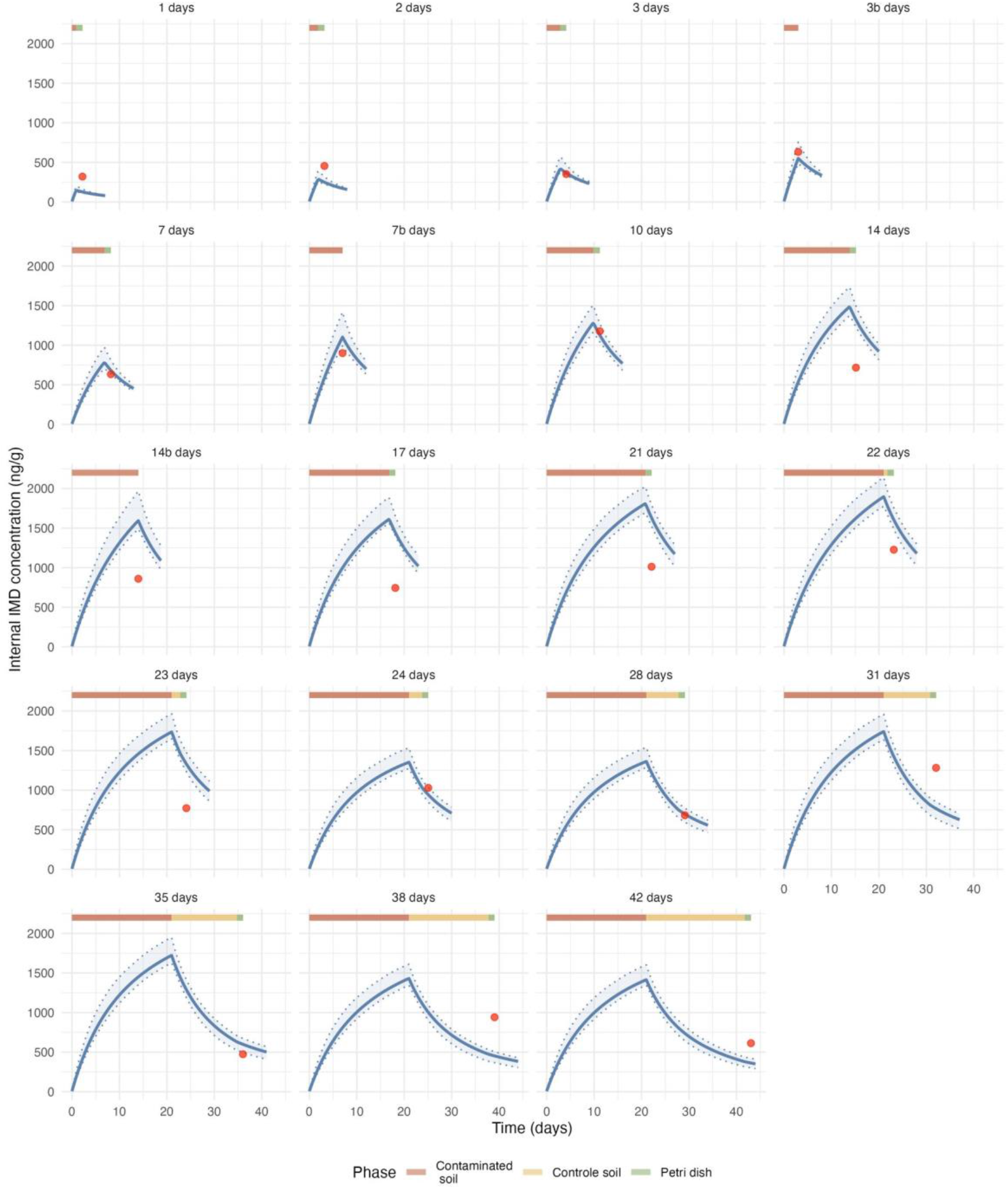
Individual estimated imidacloprid toxicokinetic in earthworms from the experiment exposed to 100 ng/g of imidacloprid. One earthworm is represented per time point (random sample). All individual trajectories are available in Supplementary information **?@fig-Art3IMDC2IndTraj1** & **?@fig-Art3IMDC2IndTraj2**. Red points represent the experimental data, the blue curves represent the estimated toxicokinetic curves and the transparent bands correspond to the credibility intervals of the model.

**Table 2:** Performance of the different toxicokinetic models for single exposure of *Aporrectodea caliginosa* to imidacloprid and epoxiconazole. The best model for each pesticide is indicated with an asterisque. C1 : One compartment model, C2 : Two-compartment model, P : Exposure through gut route in Petri dishes, N : Omission of the depuration phase in Petri dishes.

| Molecule | Model | Log-Likelihood | BIC (ΔBIC vs. best model) | Predictions within 2-folds (%) | Predictions within 5-folds (%) |
| --- | --- | --- | --- | --- | --- |
| IMD | C1 | -259.0 | 541.6 (8.7) | 86.1 | 100 |
| IMD | C2* | -251.7 | 532.9 | 90.3 | 100 |
| IMD | C2P | -276.5 | 588.2 (55.3) | 94.4 | 100 |
| IMD | C2N | -269.6 | 568.7 (35.8) | 87.5 | 100 |
| EPX | C1 | -1853.7 | 3730.8 (33776.5) | 11.3 | 56.3 |
| EPX | C1P | -1875.9 | 3781.1 (3826.8) | 42.3 | 60.6 |
| EPX | C2 | -127.9 | 285.2 (330.9) | 40.8 | 70.4 |
| EPX | C2P* | 40.5 | -45.7 | 91.5 | 100 |

**Table 3:** Estimated toxicokinetic parameters for single exposure of *Aporrectodea caliginosa* to imidacloprid and epoxiconazole (best models). Results are expressed as the modes of the distributions and their respective credibility interval (95%). The best model for each pesticide is indicated with an asterisque. C2 : Two-compartment model, P : Exposure through gut route in Petri dishes.

| | Model | $k_u$ ($\text{g}_{\text{soil}} \cdot \text{g}_{\text{worm,wwt}}^{-1} \cdot \text{d}^{-1}$) | $k_e$ ($\text{d}^{-1}$) | $k_{\text{periph}}$ ($\text{d}^{-1}$) | $\phi$ (%) | $\text{BAF}_k$ ($\text{g}_{\text{soil}} \cdot \text{g}_{\text{worm,wwt}}^{-1}$) |
| --- | --- | --- | --- | --- | --- | --- |
| IMD | C2* | 2.4 [2.1 ; 3.2] | 0.12 [0.096 ; 0.17] | 0.043 [0.025 ; 0.063] | - | 21 [18 ; 23] |
| EPX | C2P* | 1.8 [1.5 ; 2.3] | 1.8 [1.8 ; 2.0] | $1.0 \times 10^{-4}$ [$1.0 \times 10^{-4}$ ; $1.5 \times 10^{-4}$] | 67 [45 ; 98] | 0.87 [0.85 ; 1.2] |

In the case of epoxiconazole, the one-compartment model (C1) showed poor predictive performance with a clear underestimation of the steady-state and a poorly captured elimination dynamic (see Supplementary information Section S3.7). The two-compartment model (C2) slightly improved the fit but the most probable model is the two-compartment model with an exposure through the gut route in Petri dishes (C2P). All predictions fell within the 5-fold change region, with 91.5% within the 2-fold change region (Figure 2), indicating excellent predictive performance for internal concentrations in individual earthworms, with individual trajectories closely matching observed data points (Figure 5). This improvement of model predictions is reflected in a much lower BIC (Table 2). It should be noted, however, that the model still tends to overestimate internal concentrations during the second elimination phase (see Supplementary information Figure S3.24). The estimated toxicokinetic parameters of all tested models are summarized in Table 3.

**Figure 5:**
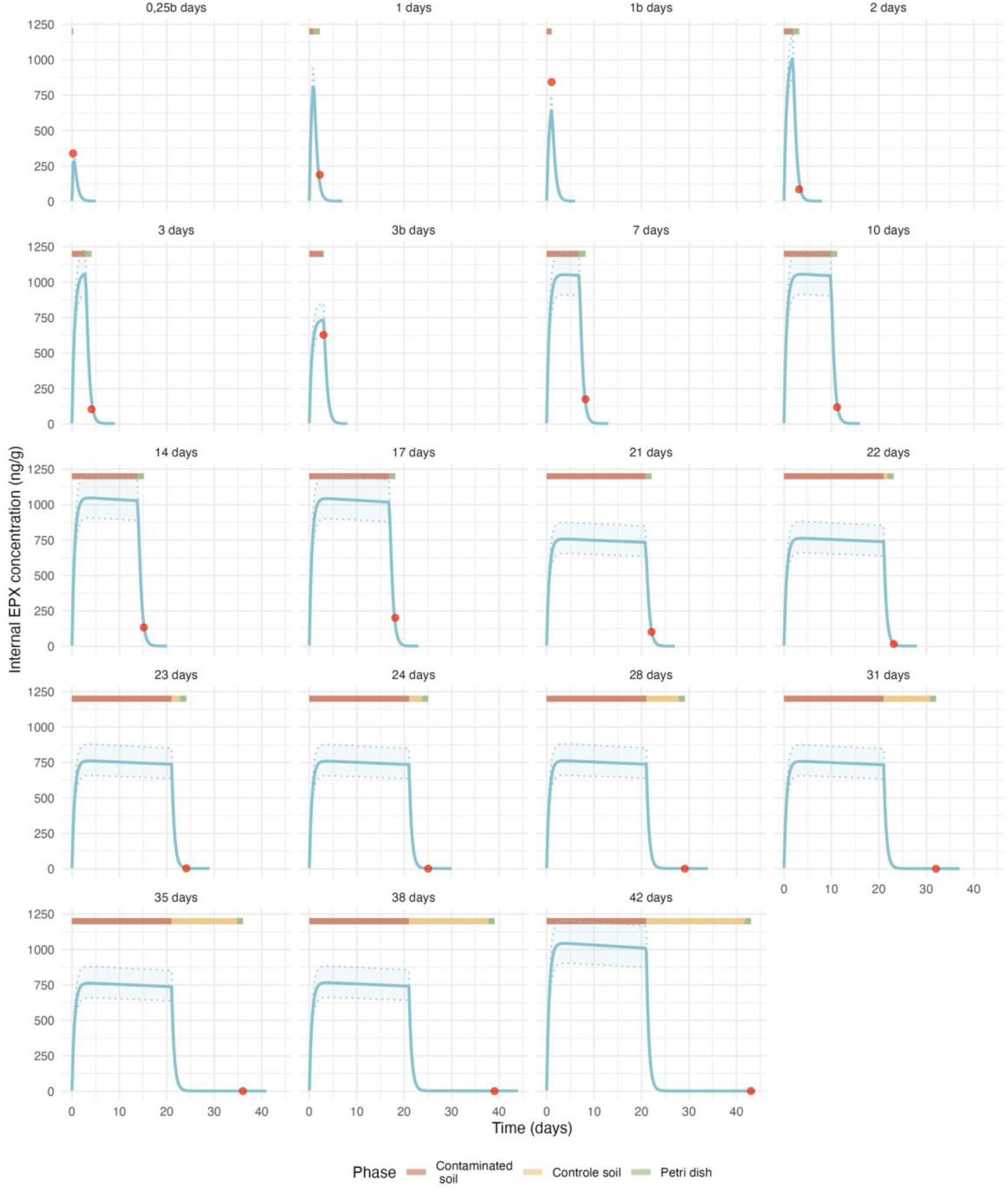
Individual estimated epoxiconazole toxicokinetic in earthworms from the experiment exposed to 1000 ng/g of epoxiconazole. One earthworm is represented per time point (random sample). All individual trajectories are available in Supplementary information **?@fig-Art3EPXC2PIndTraj1** & **?@fig-Art3EPXC2PIndTraj2**. Red points represent the experimental data, the blue curves represent the estimated toxicokinetic curves and the transparent bands correspond to the credibility intervals of the model.

Model simulations of concentration dynamics in the average earthworm under the exposure scenario of the first toxicokinetic experiment (TK) (excluding the fasting phase used for chemical analysis during which elimination occurred) revealed distinct kinetic behaviors between the two pesticides (Figure 3). Epoxiconazole exhibited rapid uptake and elimination kinetics, while imidacloprid showed considerably slower uptake and elimination processes. Correspondingly, the bioaccumulation factor (BAF) was substantially higher for imidacloprid (21 g_soil_⋅g_worm,wwt_^−1^) compared to epoxiconazole (0.87 g_soil_⋅g_worm,wwt_^−1^), reflecting its greater tendency to accumulate in earthworm tissues and slower clearance from the organism.

### 3.3 Bioaccumulation in earthworms exposed to the mixture

Single substances were also tested at two concentrations within the mixture experiment and served to evaluate whether the toxicokinetic models calibrated on single-compound exposures remained valid under mixture conditions (Figure 6). Model predictions were compared with internal concentrations measured in earthworms exposed to different mixture doses and concentration ratios. Overall, the single-substance models adequately described the mixture data, providing no evidence that co-exposure altered the toxicokinetics of either compound.

**Figure 6:**
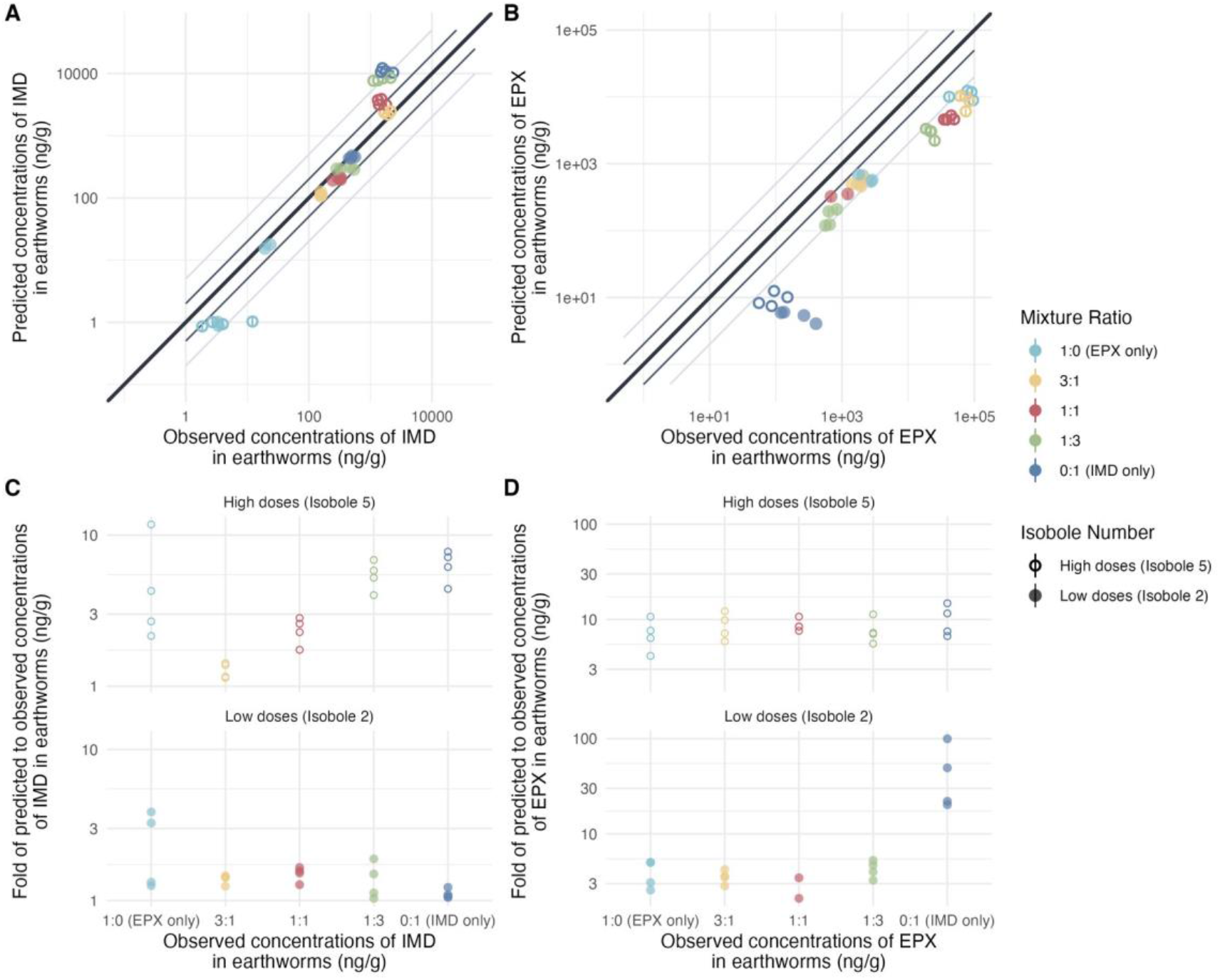
Predicted versus observed values and their respective folds for imidacloprid (two-compartment model, A & C) and epoxiconazole (two-compartment model, exposure through the gut route in Petri dishes, B & D) internal concentrations in earthworms from the mixture experiment. Points are colored by mixture ratios and the shape depends on the isobole lines (level of Toxic Unit). Grey and light-grey lines represent the 2-folds and 5-fold changes, respectively. The bold black line represents the identity line.

For both pesticides, internal concentrations measured at the lower exposure level were well predicted by the corresponding single-substance models. At higher concentrations, however, some deviations were observed. For imidacloprid, internal concentrations appeared to reach a plateau in earthworm tissues, with a mean internal concentration of 1676 ng/g, suggesting a saturation process not accounted for by the model. In contrast, epoxiconazole internal concentrations were systematically underestimated, with a mean predicted-to-observed ratio of 0.34. Nevertheless, these discrepancies were consistent across all mixture treatments, and no systematic variation with mixture ratio was observed.

Additionally, the residual concentration of epoxiconazole measured in earthworms exposed only to imidacloprid was not captured by the model, which predicted complete elimination in the absence of external exposure. This residual concentration may reflect a low background burden of epoxiconazole retained in earthworm tissues following long-term laboratory culture.

## 4 Discussion

### 4.1 Two pesticides with very different toxicokinetics

The two tested pesticides exhibited markedly different toxicokinetic behaviors in earthworms. Both pesticides showed a similar uptake but epoxiconazole has a much faster elimination than imidacloprid (*k_u_* = 1.8 vs 2.4 g_soil_⋅g_worm,wwt_^−1^⋅ d^−1^; *k_e_* = 1.8 vs 0.12 d^−1^), resulting in a much lower bioaccumulation factor (BAF = 0.9 vs 21 g_soil_⋅g_worm_^−1^). This indicates that epoxiconazole reaches equilibrium rapidly and is efficiently eliminated, whereas imidacloprid accumulates progressively over time due to its slow depuration.

Surprisingly, these kinetic patterns are opposite to what would be expected from their physicochemical properties.

Imidacloprid, although more hydrophilic (log*K_ow_* = 0.57; log *K_oc_* = 2.3), showed slow elimination and high internal concentrations. Its low log*K_ow_* indicates strong hydrophilicity, and its moderate log *K_oc_* suggests limited sorption to soil organic matter and high mobility in soil. This could reflect binding to internal aqueous compartments or specific tissue affinity (e.g. for nicotinic acetylcholine receptors or proteins), leading to retention despite low lipophilicity. Comparable kinetic profiles have been reported in aquatic invertebrates for imidacloprid: in *Gammarus pulex*, *k_u_* and *k_e_* estimates range from 1.96 - 5.21 L⋅kg^−1^⋅ d^−1^ and 0.12-0.27 d^−1^ respectively across studies (Ashauer et al., 2010; Huang et al., 2021; Mangold-Döring et al., 2022), and in *Centroptilum dipterum*, an even slower elimination (*k_e_* = 0.04 d^−1^) was associated with a higher BCF of 70.1 L⋅kg^−1^ (Huang et al., 2021), consistent with the slow depuration observed in earthworms. Overall, these findings confirm that bioaccumulation potential cannot be reliably predicted from log *K_ow_* alone, and that compound- and organism-specific processes must be considered (Arnot & Gobas, 2006; Li et al., 2024).

In contrast, epoxiconazole is considerably more hydrophobic (log*K_ow_* = 3.3; log*K_oc_* = 3.3) and moderately adsorbed to soil organic matter, with low mobility in soils. Based on hydrophobicity alone, stronger partitioning into lipid-rich tissues, slower elimination, and higher bioaccumulation would be expected. Instead, epoxiconazole was rapidly eliminated and only weakly accumulated. This pattern of fast toxicokinetics despite relatively high hydrophobicity appears consistent across organisms and azole compounds: in *G. pulex*, Rösch et al. (2016) reported very rapid uptake and elimination of epoxiconazole (*k_u_* = 217 L⋅kg^−1^⋅ d^−1^; *k_e_* = 6.66 d^−1^) with a more moderate BAF of 32.5 L⋅kg^−1^, and similarly fast elimination has been reported for other azole fungicides such as imazalil (log*K_ow_* = 3.8; *k_e_* = 360 d^−1^; BAF = 22.85 L⋅kg^−1^; (Kuhlmann et al., 2019)) and tebuconazole in zebrafish (*Danio rerio*) (log*K_ow_* = 3.7; BAF = 38.8 L⋅kg^−1^ (Andreu-Sánchez et al., 2012)).

The two-compartment structure retained for both compounds (C2) suggests that distribution processes within the organism contribute to the observed kinetics (*Δ*BIC_IMD_ (C2 vs. C1) = -8.7 ; *Δ*BIC_EPX_ (C2 vs. C1) = -3445.6 ; Table 2). For hydrophobic compounds in terrestrial systems, biphasic elimination has frequently been reported and linked to soil-related processes (Belfroid & Sijm, 1998). The initial rapid phase may be influenced by interactions with soil organic matter leading to a change in the exposure, while a slower phase can reflect redistribution within the organism..

In contrast, the inclusion of an additional exposure pathway through residual gut content during the depuration phase was only supported for epoxiconazole (C2P), and not for imidacloprid (*Δ*BIC_EPX_ (C2P vs. C2) = -330.9 ; Table 2). This difference is consistent with physicochemical expectations: exposure via gut content is generally considered negligible for hydrophilic compounds such as imidacloprid, but may represent a significant fraction of total uptake for hydrophobic substances (log *K_ow_* > 4.5), with reported contributions ranging from 15 to 40% (Belfroid et al., 1995; Jager et al., 2003). Although epoxiconazole is only moderately hydrophobic (log *K_ow_* = 3.3), the model results suggest that this pathway may still contribute to its overall kinetics.

The underlying mechanisms therefore likely differ between the two compounds. For imidacloprid, the two-compartment structure primarily reflects internal distribution processes. For epoxiconazole, in addition to internal distribution, exposure through residual gut content may contribute to the observed dynamics, potentially through adsorption of the compound to gut surfaces with continued desorption of contaminated particles. Reuptake from the surrounding soil is unlikely in the present study due to the large soil volume relative to earthworm biomass. Additionally, the presence of an aqueous stagnant layer surrounding the organism may act as a diffusion barrier; adsorption of the compound onto soil particles could facilitate transport across this layer, thereby influencing elimination kinetics. More broadly, it suggests that, in soil-fauna studies, evaluating a two-compartment model may also be relevant for hydrophilic substances.

Together, these results highlight that hydrophobicity alone is insufficient to predict toxicokinetic behavior in earthworms and that interactions among soil properties, organism physiology, and compound-specific processes jointly determine uptake and elimination dynamics.

### 4.2 Importance of a well described exposure scenario in toxicokinetic modeling

Our results highlight the critical importance of explicitly describing and integrating the experimental exposure scenario into toxicokinetic (TK) models, especially for rapid TK compounds. In standard bioaccumulation tests following OECD TG 317 (OECD, 2010), earthworms (*Eisenia* sp.) are allowed to purge their gut overnight on moist filter paper in covered Petri dishes prior to chemical analysis. However, the guideline does not specify how this depuration phase should be handled in TK modeling, nor how it may influence the estimation of bioaccumulation factors (BAF). In practice, BAF_ss_ is calculated as the ratio between mean organism and soil concentrations at steady state, while BAF_k_ is derived from the ratio *k_u_*/*k_e_* when steady-state is not reached. The implicit assumption is that the overnight depuration step does not substantially alter internal concentrations.

Our study demonstrates that this assumption may not hold for substances with rapid toxicokinetics.

For epoxiconazole, which exhibited fast uptake and fast elimination, steady state was effectively reached within the first day of exposure. If BAF*_ss_* were calculated strictly according to OECD TG 317 (i.e., using internal concentrations measured after overnight depuration), the resulting value would be 0.14 g_soil_⋅g_worm,wwt_^−1^. In contrast, the TK model, explicitly accounting for internal dynamics and the depuration phase (C2P), estimated a BAF of 0.9 g_soil_⋅g_worm,wwt_^−^1. This discrepancy illustrates how a 24-hour depuration step can strongly bias bioaccumulation estimates for rapidly eliminated compounds. In such cases, the depuration phase is not negligible relative to the characteristic time constants of uptake and elimination and must be explicitly represented in the model.

For imidacloprid, the influence of the Petri dish phase was less pronounced. Comparing the C2 model with the same structure assuming no net exchange during this period (C2N), which effectively neglects the Petri dish phase, showed that explicitly accounting for this phase provided a better description of the data (*Δ*BIC(C2 vs. C2N) = -35.8). It resulted in similar estimated BAF values between the two models (21 vs. 18 g_soil_⋅g_worm,wwt_^−1^), with overlapping confidence intervals (C2: 21 [18 ; 23]; C2N: 18 [16 ; 21]), indicating no significant difference. More information about the C2N model is available in the Supplementary information Section S3.4).

A review of recent studies indicates that the overnight gut-purging step is routinely implemented (Del Puerto et al., 2025; Heinrich et al., 2025; Lotufo et al., 2025), yet it is rarely incorporated into TK model structures. In earlier work such as Jager (1998), lower-than-expected BCF values were reported for highly lipophilic compounds (e.g., PCBs), which could plausibly be influenced by unaccounted elimination during depuration. Although this explanation would require case-specific verification, it illustrates the broader issue that depuration phases may systematically bias kinetic parameter estimation if not modeled explicitly.

These considerations suggest that, for substances suspected to exhibit rapid uptake or elimination, sampling time points shorter than 24 hours should be included in both uptake and elimination phases. In addition, alternative designs, such as immediately sampling earthworms and manually massaging them to void gut contents rather than allowing a 24-hour depuration on Petri dishes, could help avoid the bias in internal concentrations introduced by the depuration phase. The additional individuals that were only massaged and were not subjected to the depuration phase on Petri dishes further improves the estimation of steady-state concentrations and BAF. This is especially important for compounds such as epoxiconazole, characterized by rapid uptake and elimination kinetics, for which substantial elimination already occurs during the 24-hour depuration period. In the original design, the first sampling point occurred after one day of exposure followed by one day of depuration, thereby missing the early phase of apparent uptake. As a result, the apparent uptake rate cannot be estimated with sufficient precision. Introducing an earlier sampling time (e.g., 6 hours) combined with the absence of a depuration phase allows for a more accurate characterization of the uptake phase and a more reliable estimation of steady state. This refined design likely also improves parameter estimation for imidacloprid, although to a lesser extent given its slower toxicokinetic dynamics.

Overall, our findings emphasize that the Petri dish depuration step is not a trivial procedural detail but a kinetic phase in its own right. Its duration relative to compound-specific rate constants determines whether it is negligible or whether it substantially alters apparent bioaccumulation metrics. Explicit integration of this phase into TK modeling should therefore become standard practice when interpreting earthworm bioaccumulation data.

### 4.3 No interaction detected in the mixture at the toxicokinetic level

The mixture experiment did not reveal any detectable interaction between epoxiconazole and imidacloprid at the toxicokinetic level. No consistent trend in internal concentrations was observed across mixture ratios, and the hypothesis that epoxiconazole could limit the elimination of imidacloprid was therefore not supported under our experimental conditions. If such an interaction exists, it is likely to be small relative to the intrinsic variability of the system. These findings are consistent with the toxicodynamic results, where only a weak synergistic interaction was detected (Gollot et al., 2026b), suggesting that interactions between epoxiconazole and imidacloprid are limited at both TK and TD levels under the tested conditions.

One plausible explanation for the absence of a strong toxicokinetic interaction is that imidacloprid elimination in earthworms may not rely predominantly on cytochrome P450-mediated metabolism. While metabolic inhibition via P450 interference is a well-documented mechanism in insects and some aquatic invertebrates, earthworms possess alternative detoxification pathways that could facilitate the clearance of imidacloprid independently of P450 activity. These include glutathione-S-transferase-mediated conjugation, carboxylesterase activity, antioxidant enzymes, and microbial metabolism in the gut, all of which may contribute to the transformation or excretion of imidacloprid (Katagi & Ose, 2015; Wang et al., 2019). Additionally, direct excretion of the parent compound through passive diffusion may also play a significant role, particularly given imidacloprid’s moderate hydrophilicity, which favors movement in aqueous compartments. Taken together, these alternative elimination routes could buffer potential metabolic interactions with co-occurring substances such as epoxiconazole. Consequently, any effect of epoxiconazole on imidacloprid metabolism is likely to be minimal, consistent with the lack of detectable interaction in internal concentrations. This highlights the importance of considering species-specific detoxification mechanisms when extrapolating metabolic interactions across taxonomically distant invertebrates.

### 4.4 Dependency of the BAF with the exposure concentration

Beyond interaction testing, the mixture experiment addressed an important limitation of the single-substance TK experiments, in which only one exposure concentration per compound was tested. Although this design choice is common in TK studies due to their experimental intensity, it precludes evaluation of potential concentration dependency of kinetic parameters or bioaccumulation factors. The mixture experiment, by including ten exposure levels per substance, provides additional insight into this aspect and suggests that concentration-dependent processes occur for both pesticides at higher exposure levels, although only a single sampling time point (28 days) was available.

For imidacloprid, internal concentrations measured at high exposure levels indicate the onset of a saturation process. Several non-exclusive mechanisms could account for this pattern. At elevated concentrations, behavioral changes may reduce feeding activity and thereby limit uptake. For instance, reduced cast production has been reported in *A. caliginosa* and *Lumbricus terrestris* at imidacloprid concentrations ≥ 0.66 mg⋅kg^−1^ dry soil after 7 days of exposure, while stimulation was observed at the lower concentration tested (0.2 mg⋅kg^−1^) (Dittbrenner et al., 2010, 2011). Such sublethal effects on feeding behavior could directly alter uptake rates and the concentration threshold for reduced cast production might be dependent on the soil used. Saturation of tissue binding sites at higher internal concentrations represents another plausible explanation. Depletion of the exposure medium can be excluded, as the total mass of imidacloprid accumulated in individual earthworms remained negligible relative to the total amount present in the cosm (about 10%).

For epoxiconazole, interpretation is more complex. The mixture experiment contained approximately twice as much horse dung as the single-substance experiment, resulting in a higher organic matter content. Given that epoxiconazole is moderately lipophilic (log *K_ow_* = 3.3) and has a relatively high log *K_oc_* (3.3), stronger sorption to organic matter would be expected, potentially reducing the freely dissolved fraction available for uptake. A higher log *K_oc_* generally implies reduced bioavailability to earthworms due to increased binding to soil organic matter (Del Puerto et al., 2025). However, the model underestimated internal epoxiconazole concentrations by a similar factor across all tested conditions, with no clear pattern related to mixture ratio. This suggests that the discrepancy more likely reflects a general difference in exposure conditions between experiments, rather than a mixture-driven toxicokinetic interaction. One possible explanation is that binding sites in soil organic matter became saturated at higher epoxiconazole concentrations, thereby increasing the freely available fraction. Changes in feeding behavior or ingestion of organic matter-associated residues may also contribute. At present, these mechanisms remain hypothetical and cannot be disentangled with the available data.

Concentration-dependent bioaccumulation has been reported in other soil invertebrate systems. For instance, in *Eisenia andrei* exposed to PFAS mixtures, BAFs at 1 mg⋅kg^−1^ were more than one order of magnitude lower than at 0.01 mg⋅kg^−1^, a pattern partly attributed to sublethal physiological impairment at higher concentrations (Lotufo et al., 2025). Such behavior challenges the assumption of concentration-independent BCF or BAF values for hazard classification, particularly for polar or metabolizable organic compounds.

Although the high concentrations tested here exceed median environmental levels (Table 4) and therefore have limited direct environmental relevance, they provide important mechanistic insights. In particular, these concentration-dependent patterns would likely have remained undetected if only low concentrations with no observed effect on the organisms had been tested, as commonly used in standard toxicokinetic studies. If internal concentrations are not proportional to external exposure beyond a certain threshold, then toxicodynamic studies conducted at high doses may not reflect a simple exposure-internal dose relationship. This again underscores the importance of jointly considering toxicokinetic and toxicodynamic processes when interpreting effect data.

**Table 4:** Summary of results for the detected imidacloprid and epoxiconazole residues in the Froger et al. (2023), Pelosi et al. (2021) and Silva et al. (2019) studies.

| **Molecule** | **Former application dose (EU) (Pelosi et al., 2021)** | **Source** | **Median (ng/g)** | **Q3 (ng/g)** | **Maximum (ng/g)** |
| --- | --- | --- | --- | --- | --- |
| IMD | 168 ng/g | Froger et al. (2023) | 2.9 | 5.8 | 13.8 |
|  |  | Pelosi et al. (2021) | 15.1 | - | 160 |
|  |  | Silva et al. (2019) | 20 | - | 60 |
| EPX | 153 ng/g | Froger et al. (2023) | 5.8 | 10.9 | 23.6 |
|  |  | Pelosi et al. (2021) | 34.6 | - | 283 |
|  |  | Silva et al. (2019) | 20 | - | 160 |

### 4.5 Perspectives

Some limitations of the present study should be acknowledged, while also pointing to directions for future research. First, working with soil organisms inherently involves variability related to soil properties and background contamination, which may influence bioavailability and toxicokinetic processes. In particular, soil organic matter is known to play a key role in the dynamics of hydrophobic compounds (Belfroid & Sijm, 1998).

Second, the use of total soil concentrations rather than bioavailable fractions may have introduced uncertainty in bioaccumulation estimates. Future studies incorporating measurements of porewater or water-extractable concentrations could improve the interpretation of uptake processes and help explain the concentration-dependent patterns observed. More broadly, considering changes in pesticide availability over time, such as residue aging, would also help better characterize exposure dynamics in soil systems (Šudoma et al., 2021).

Finally, metabolites were not quantified in this study. Including metabolite analysis in future work would provide a more comprehensive understanding of internal exposure and transformation processes. For chiral compounds, this could also include enantiomer-specific analyses, as stereoselective degradation and bioaccumulation may influence both exposure and toxicity (Bielská et al., 2021; Škulcová et al., 2020).

Overall, addressing these aspects and testing whether the present conclusions hold across other pesticides, soil types, and earthworm species would help improve the mechanistic understanding of pesticide behavior in soil invertebrates. From a modeling perspective, integrating both toxicokinetic and toxicodynamic components into a unified DEB-TKTD framework would further allow a more mechanistic characterization of individual substances and their mixtures, including potential concentration-dependent processes (Jager, 2015; Kooijman et al., 2009). More broadly, our results highlight the need to jointly consider experimental design, exposure characterization, and model structure when studying bioaccumulation in soil systems. Such integrated approaches are essential to refine toxicokinetic modeling and strengthen the environmental relevance of mixture risk assessment in terrestrial invertebrates.

## 5 Conclusion

This study provides a detailed assessment of the toxicokinetics of two pesticides, epoxiconazole and imidacloprid, in the earthworm *A. caliginosa*, both as single substances and in mixtures. The two compounds exhibited strikingly different kinetic behaviors: imidacloprid, despite its hydrophilic nature, accumulated slowly and was eliminated very slowly, whereas epoxiconazole, moderately lipophilic, reached equilibrium rapidly and was efficiently eliminated. These patterns highlight that bioaccumulation potential cannot be reliably predicted from log*K_ow_* or log*K_oc_* alone, and that compound-specific physiological and soil-organism interactions play a critical role.

The use of two-compartment toxicokinetic models allowed a more mechanistic representation of internal distribution and redistribution dynamics, capturing biphasic uptake and elimination for both substances. In particular, explicitly modeling the depuration phase on Petri dishes proved essential for accurate parameter estimation, especially for epoxiconazole. The mixture experiment revealed no detectable toxicokinetic interactions between epoxiconazole and imidacloprid, indicating that single-substance models can reliably describe uptake and elimination in this context.

Additionally, the study emphasizes the importance of considering concentration-dependent processes. At high exposure levels, both compounds showed signs of saturation, whether at the tissue level or in relation to soil organic matter binding, underscoring that internal concentrations may not scale linearly with external exposure. This finding questions the use of single, concentration-independent bioaccumulation factors for hazard assessment and highlights the value of integrated TKTD approaches.

Finally, our work demonstrates the importance of well-characterized experimental exposure scenarios and careful handling of soil organisms in toxicokinetic studies. Incorporating short-term depuration, gut-content dynamics, and variability in soil properties improves model reliability and the interpretation of bioaccumulation data. Overall, the combined use of mechanistic toxicokinetic models and controlled experimental designs provides a robust framework to understand and predict the uptake, distribution, and elimination of chemicals in soil invertebrates, with implications for both single substances and mixtures in environmental risk assessment.

## Data availability statement

The data produced in this study will be available in on data.gouv.fr after submission (Gollot et al., 2026a). All R scripts used in this study are publicly available on GitHub (https://github.com/LisaGllt/Ew-Mix-TK) with an accompanying website providing comprehensive documentation and in-depth explanations (https://lisagllt.github.io/Ew-Mix-TK).

## Author contribution statement

L. Gollot : Conceptualization, Data curation, Formal Analysis, Investigation, Methodology, Software, Validation, Visualization, Writing – Original Draft Preparation.

J. Faburé : Funding Acquisition, Methodology, Resources, Supervision, Writing – Review & Editing.

G. Delarue : Data curation, Formal Analysis, Investigation, Methodology, Validation, Writing – Review & Editing.

L. Frattaroli : Data curation, Investigation, Methodology, Writing – Review & Editing.

S. Nélieu : Data curation, Investigation, Methodology, Validation, Supervision, Writing – Review & Editing.

R. Royauté : Conceptualization, Funding Acquisition, Methodology, Project Administration, Resources, Supervision, Writing – Review & Editing.

R. Beaudouin : Formal Analysis, Funding Acquisition, Methodology, Resources, Supervision, Writing – Review & Editing.

## Disclaimer

The authors declare that they have no known competing financial interests or personal relationships that could have appeared to influence the work reported in the present study.

## Acknowledgments

The authors thank M. Collombel, V. Le Bars, C. A. T. Sarr, J. Rouyer, A. Bamière and V. Etievant for their technical support. The authors also thank for chemical analysis management and developments Marjolaine Deschamps, Thinhinane Hamitouche and Louis Genain.

