## Supplementary Information for "Toxicokinetics of a Pesticide Mixture in Earthworms Reveal Concentration-Dependent Bioaccumulation and Limited Interactions"

### 1 Earthworm culture

#### 1.1 Breeding and experiment soil

The soil used in the culture of earthworms is collected from the top 10 cm of a meadow in Versailles called “Les Closeaux”. It is then dried before being milled at $< 2$ mm.

Table 1.1: **Main characteristics of the breeding soil from Versailles.** Table from Bart et al. (2017).

| Characteristics | Value |
|---|---|
| Clay ( $< 2$ µm, g/kg) | 226.0 |
| Fine silt (2-20 µm, g/kg) | 174.1 |
| Coarse silt (20-50 µm, g/kg) | 298.9 |
| Fine sand (50-200 µm, g/kg) | 239.1 |
| Coarse sand (200-2000 µm, g/kg) | 47.9 |
| $CaCO_3$ total (g/kg) | 23.3 |
| Organic matter (g/kg) | 32.6 |
| $P_2O_5$ (g/kg) | 0.1 |
| Organic carbon (g/kg) | 18.9 |
| Total nitrogen (N) (g/kg) | 1.5 |
| C/N | 12.5 |
| pH | 7.5 |
| Total $Cu$ (mg/kg) | 25.2 |

#### 1.2 DNA barcoding

The phylogenetic tree was realised thanks to maximum likelyhood bootstrap method (1000 simulations) with the Tamura-Nei model (nucleotides). Bootstrap frequency is indicated on each branch.

*Aporrectodea caliginosa* is a cryptic species complex comprising three lineages. Our earthworm culture is known to include individuals from two closely related lineages (2 and 3). While these lineages could potentially exhibit differences in sensitivity, previous research on *Gammarus roeselii* has shown that tolerance within cryptic species complexes is not always lineage dependent (Kabus et al., 2024).

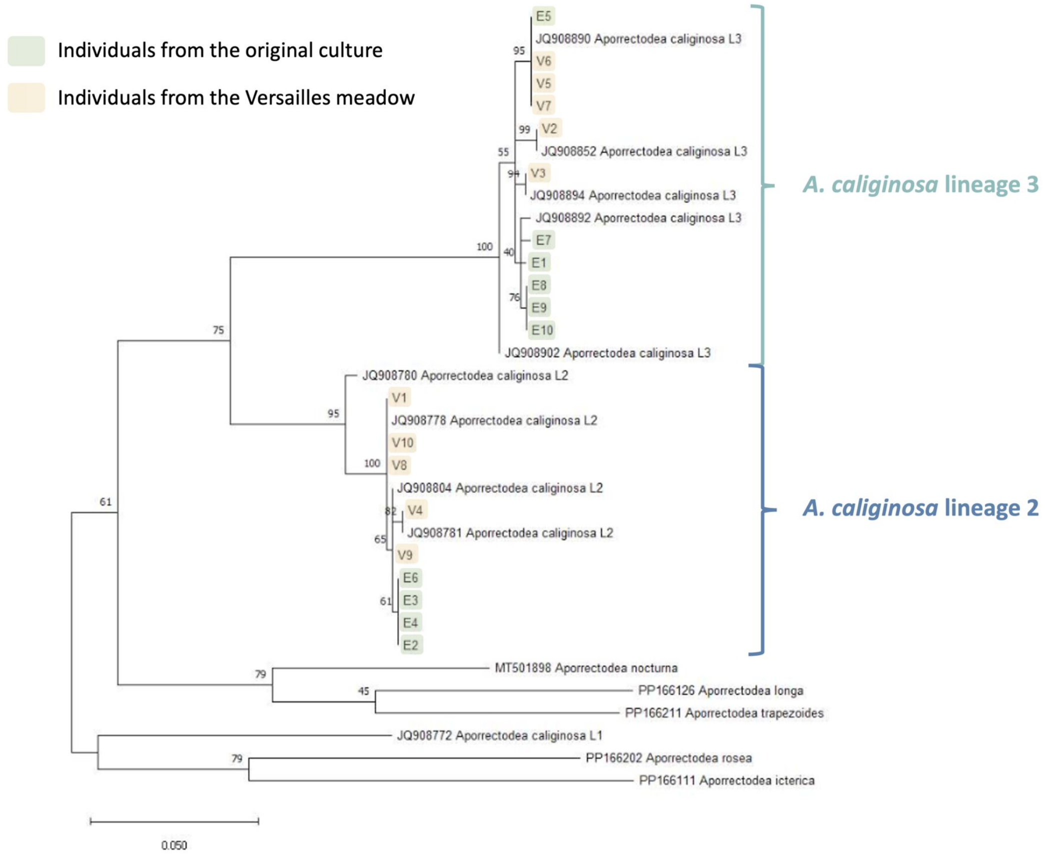

Figure 1.1: Results of the DNA barcoding of 10 individuals from the culture (highlighted in yellow)

### 2 Experiments

#### 2.1 Single substances experiments

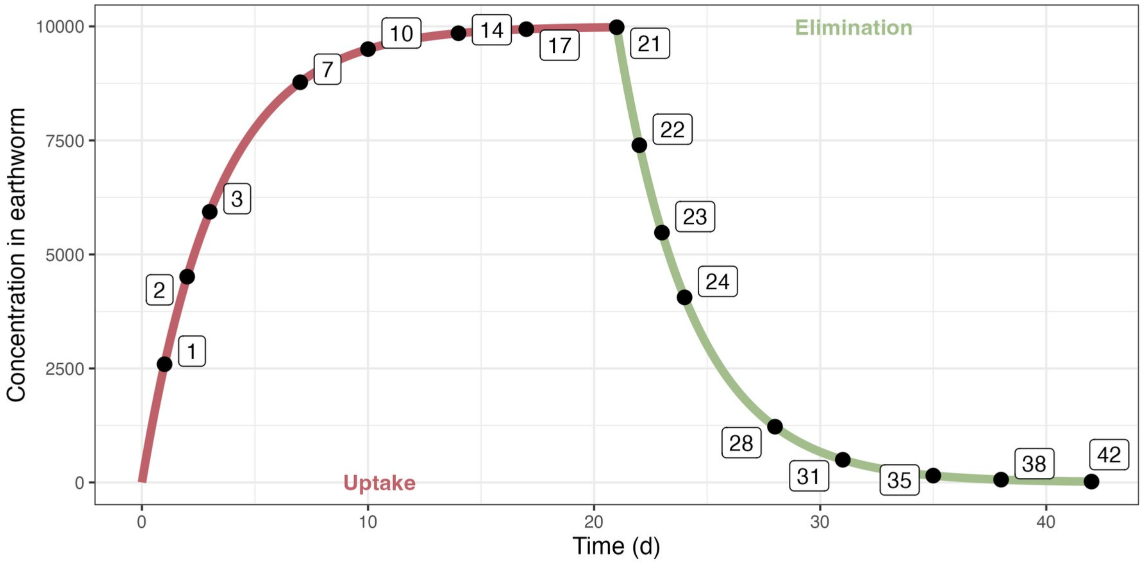

Figure 2.1: Design of the initial toxicokinetic experiments

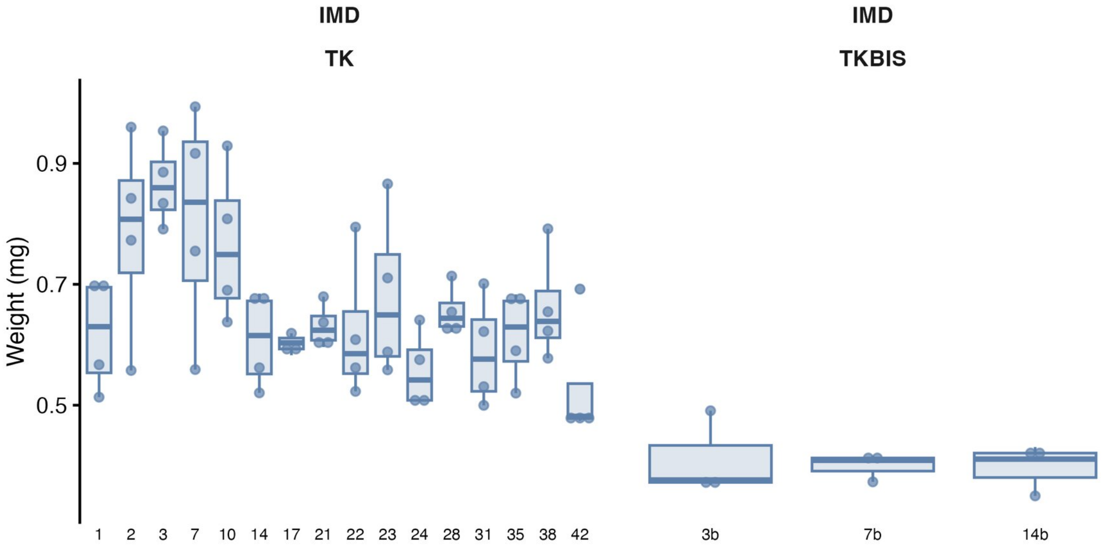

Figure 2.2: Initial weights (with gut content) of earthworms for the IMD experiment.

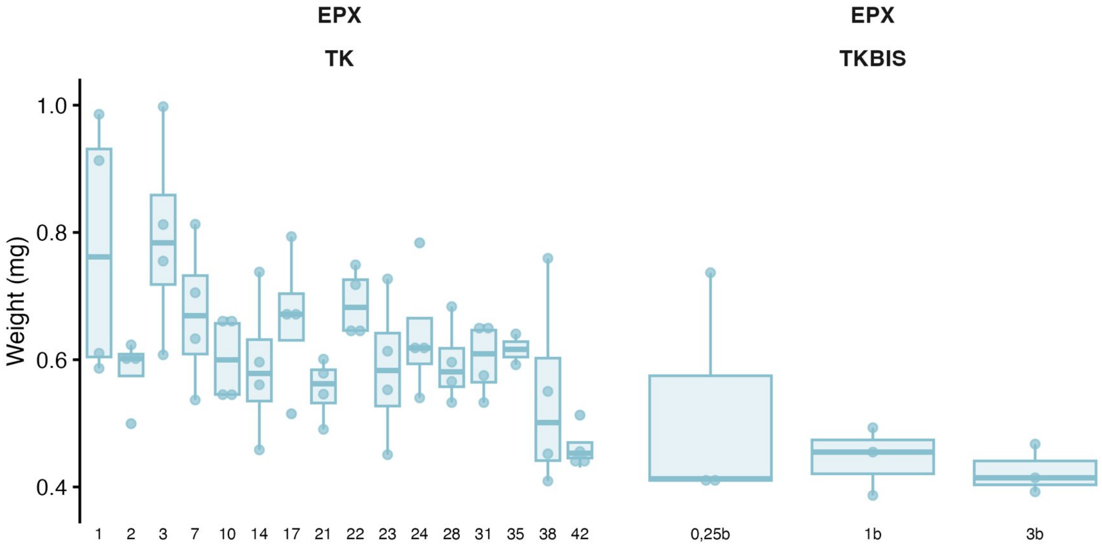

Figure 2.3: Initial weights (with gut content) of earthworms for the EPX experiment.

Table 2.1: **Initial weight of earthworms (with gut content, mg) in the toxicokinetic experiments.** The letter "b" indicates earthworms used with only a massage before chemical analysis.

| Molecule | Min | 1st Q | Mean | Median | 3rd Q | Max | $n$ |
|---|---|---|---|---|---|---|---|
| IMD (TK) | 474.0 | 565.7 | 662.5 | 635.1 | 711.3 | 993.7 | 64 |
| IMD (TK b) | 350.0 | 373.2 | 402.8 | 408.8 | 416.5 | 490.8 | 9 |
| EPX (TK) | 409.3 | 544.7 | 622.8 | 605.8 | 672.6 | 997.7 | 62 |
| EPX (TK b) | 386.7 | 408.0 | 462.9 | 414.4 | 467.4 | 736.7 | 9 |

#### 2.2 Link between weight with and without gut content

Since experimental data include weights with and without gut content, we established a quantitative relationship between them to allow conversion and ensure consistency across datasets. The relationship is made with the data from the first toxicokinetic experiments with the weight of the earthworm before and after the 24h in Petri dishes (and massage if needed). We make the assumption that the variation in body weight without gut content is negligeable during the 24 hours. The results are shown Figure 2.4.

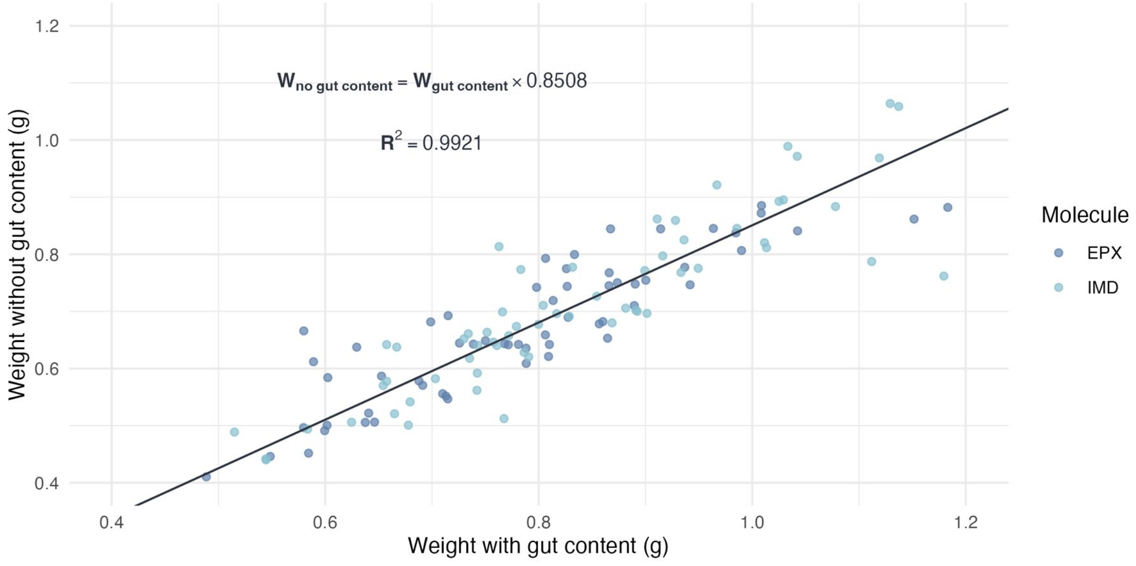

Figure 2.4: Relationship between weight with and without gut content.

#### 2.3 Depuration kinetic in Pétri dishes (GUT experiment)

Depuration of the earthworm gut content was modeled as a first-order decay (Equation 2.1), informed by a small supporting experiment: ten earthworms were kept individually in 200 g of Closeaux soil with 3 g of horse dung for one week, then transferred to Petri dishes lined with moist filter paper and weighed at regular intervals. After 32 hours, the worms were gently massaged to void the remaining gut contents and weighed again, providing data to estimate the gut-clearance rate.

$$\frac{dP_{Weight}}{dt} = -\frac{\ln 2}{DT_{50,\mathrm{gut}}} \left(P_{Weight} - P^{\infty}_{Weight}\right) \tag{2.1}$$

$$P_{Weight} = P^{\infty}_{Weight} + \left(100 - P^{\infty}_{Weight}\right) \exp\left(-\frac{\ln 2}{DT_{50,\mathrm{gut}}} \times t\right)$$

With :

- $t$ : Time (h)
- $P_{Weight}$ : Percentage of initial weight (%, 0-100)
- $P_{Weight}^{\infty}$ : Percentage of initial weight when gut is fully emptied (%)
- $DT_{50,\text{gut}}$ : Half-life of the gut content (h)

For the model, we consider that weights measured after massage are weights measured after a significant amount of time has passed. Therefore, the corresponding percentages of initial weight inform $P_{Weight}^{\infty}$.

The estimated value for $DT_{50,\text{gut}}$ and $P_{Weight}^{\infty}$ are 4.37 [3.11 ; 7.32] hours and 86.2 [84.6 ; 87.9] %, respectively. $P_{Weight}^{\infty}$ estimated here is consistent with the one estimated in Section 2.2.

$P_{Weight}^{\infty}$ estimated here is consistent with the one estimated in Section 2.2.

Model evaluation is presented Figure 2.5 and Figure 2.6.

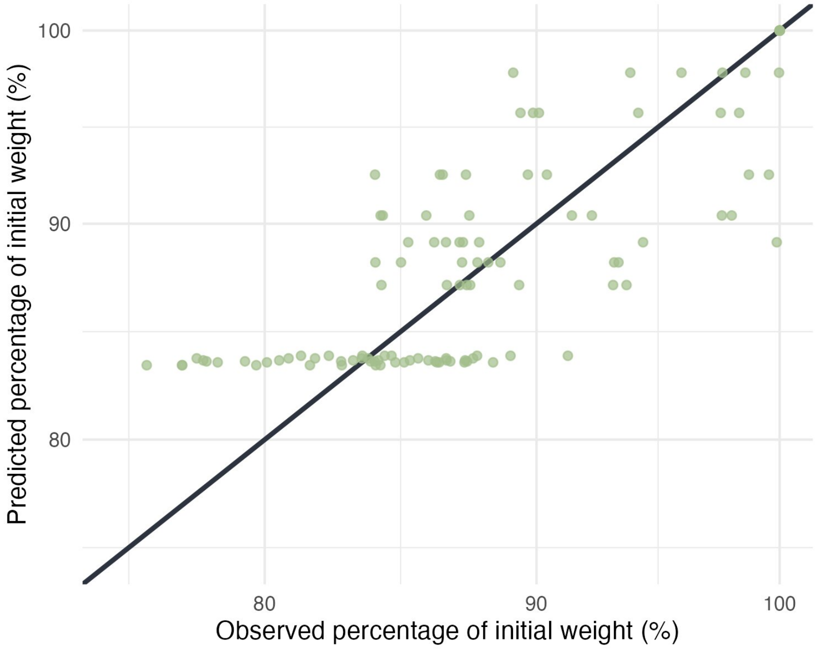

Figure 2.5: Predicted vs observed percentage of initial weight. All data points fall within fold 2.

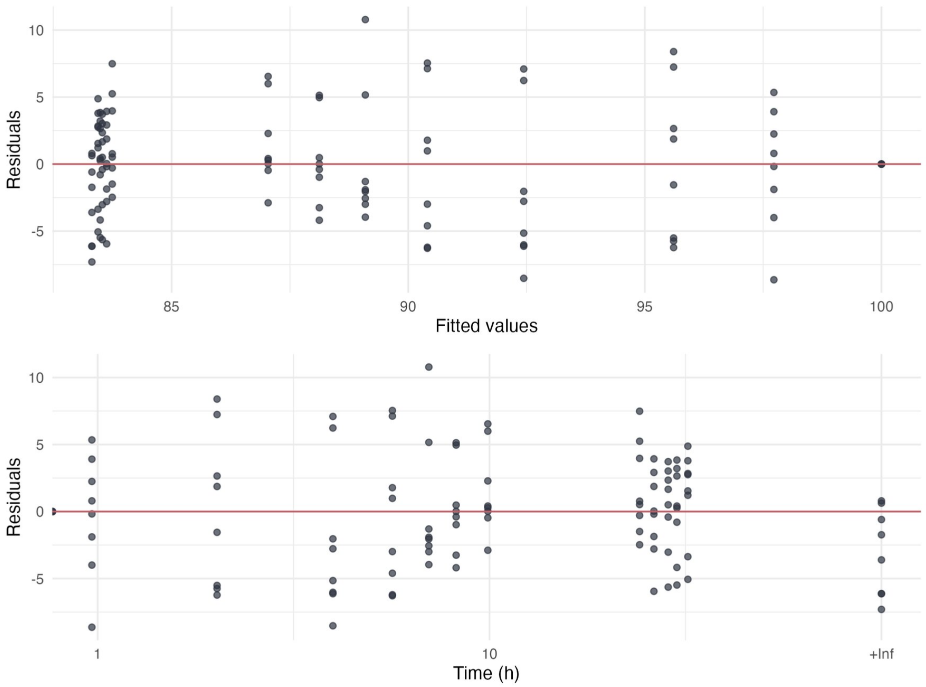

Figure 2.6: Residuals of the model for earthworm depuration.

#### 2.4 Mixture experiment

Ratio are labeled with letters : E : 1:0 (EPX only) ; F : 3:1 ; G : 1:1 ; H : 1:3 ; I : 0:1 (IMD only)

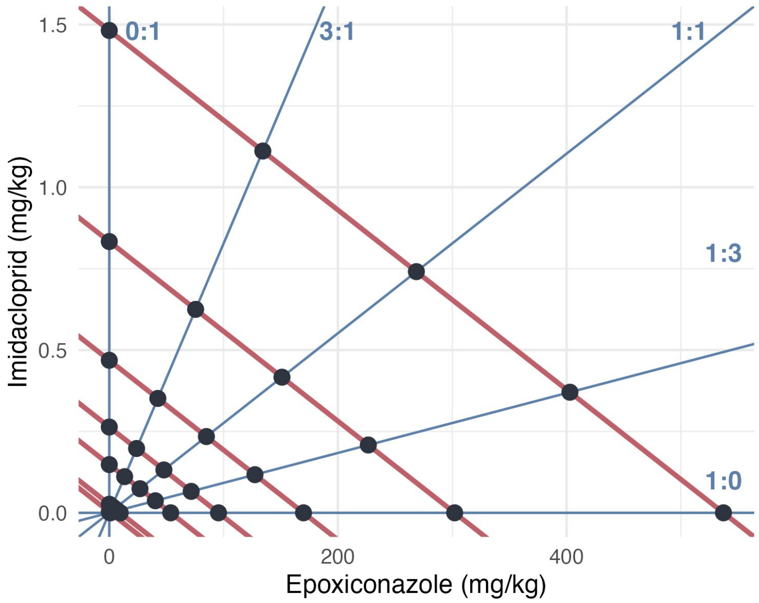

Figure 2.7: Experimental design of the mixture experiment (MIX). Blue lines represent the ratio if TU tested and the red lines correspond to effect isoboles under concentration addition hypothesis.

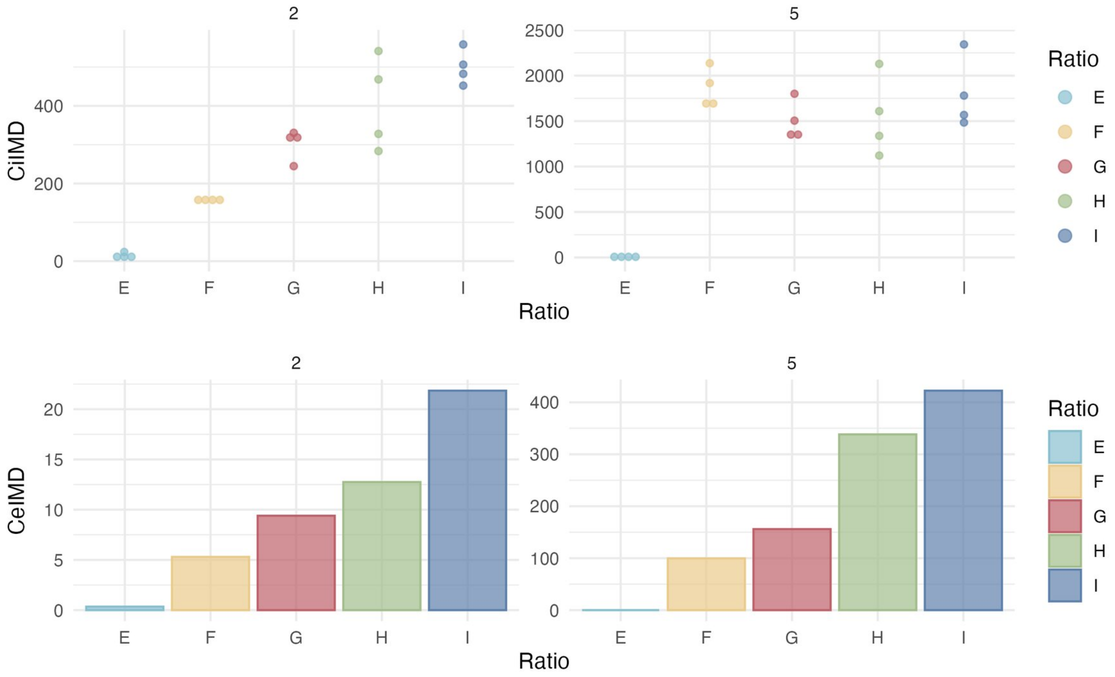

Figure 2.8: Results of the chemical analysis of internal concentrations in IMD of earthworms exposed to the mixture.

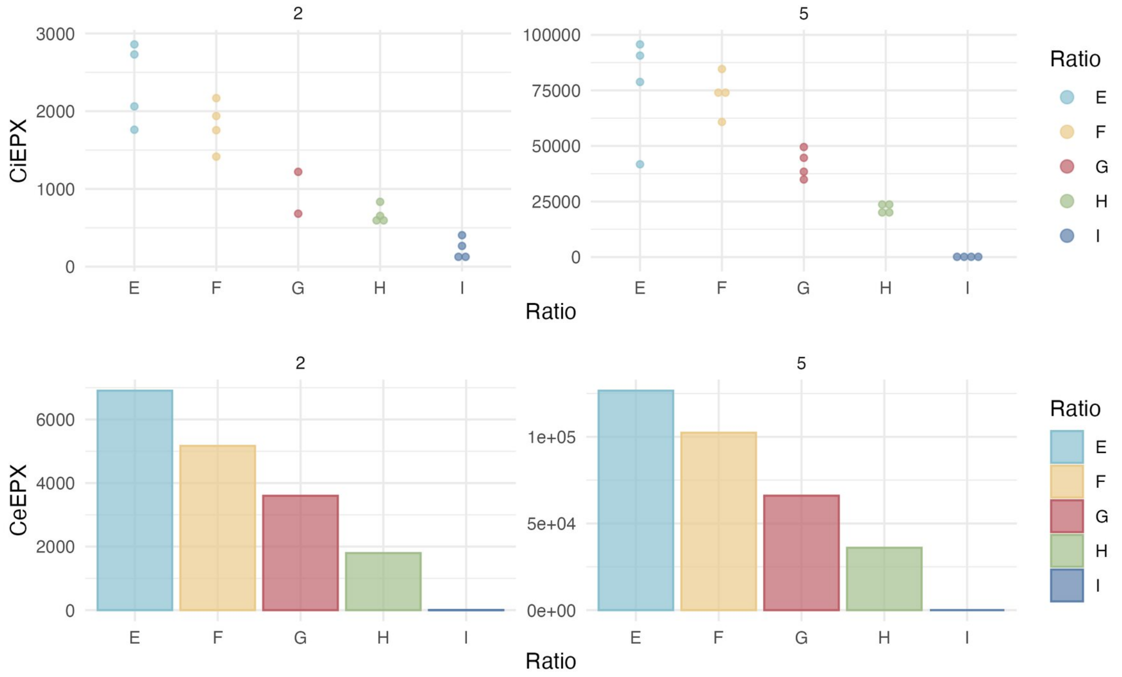

Figure 2.9: Results of the chemical analysis of internal concentrations in EPX of earthworms exposed to the mixture.

#### 2.5 Chemical analysis

##### 2.5.1 Chemicals and reagents

The two active substances IMD and EPX used as analytical standard were the same as mentioned above. Their deuterated equivalent epoxiconazole-d4 and imidacloprid-d4, used as internal standards, came from Cluzeau-Info-Labo. They were all solubilized in acetonitrile.

Acetonitrile (quality HPLC-MS), formic and acetic acids (analytical purity) were provided by Carlo Erba. The purified water called UP water, presenting a resistivity > 18.2 MΩ.cm and a total organic carbon content < 4 ppb, was produces by a Purelab Chorus I station. The QuEChERS extraction salts (4 g $MgSO_4$ + 1 g sodium acetate, Chromabond mix II) and the d-SPE sorbents (150 mg $MgSO_4$ + 50 mg C18 + 50 mg PSA, Chromabond QuEChERS mix XIX) came from Macherey-Nagel.

##### 2.5.2 Preparation of soil samples

After freeze-drying, 5 g of dry soil was weighed in a 50 mL Falcon® tube, and humidified by 1 mL of UP water to restore its sorption capabilities. After 2 h, the soil was spiked with 50 µL of a deuterated internal standards solution and kept overnight away from light and at 4°C to allow sorption to happen.

For extraction, 6 mL of water and 7 mL of acetonitrile acidified by 1% acetic acid were successively added (shaking by 5 s vortex after each addition). Then, after addition of the QuEChERS salts, the tubes were vigorously shaked, vortexed 5 s, shaked 15 min at 300 rpm on an orbital shaking table and submitted to 5 min ultra-sounds in a water bath. After centrifugation 10 min at 2095 g and 20°C, the acetonitrile upper layer was collected. Then, 1500 µL of extract were added in an Eppendorf containing the d-SPE sorbent. The Eppendorf was hand-shaked, vortexed 5 s, centrifuged 10 min at 12100 g and 400 µL of the purified extract was placed in an injection vial. The 400 µL samples were evaporated under nitrogen flow and reconstructed by 60 µL of acetonitrile and then 1140 µL of UP water acidified by 0.1% acetic acid, each liquid addition being followed by 5 s vortex. The samples were diluted if necessary (by a 50 to 200 factor). Preparation of earthworm samples

The frozen earthworms were cutted with a scalpel and 300 mg were weighed in FastPrep® microtubes. A ceramic ball and 1 mL of UP water were introduced in the microtubes and samples were crushed in a FastPrep equipment during two cycles of 20 s at 6 m/s. The crushed samples were transferred in a 50 mL Falcon® tube; the microtubes and the ceramic balls were rinsed with 5 mL UP water and this water was collected and added to the 50 mL tubes. The deuterated internal standards were then added and allowed to establish interactions with the matrix during 1 h.

For extraction, 5 mL of acetonitrile acidified by 1% formic acid was added, as well as the QuEChERS salts. The tubes were vigorously shaked, vortexed 5 s, and shaked 10 min at 300 rpm on an orbital shaking table. After centrifugation 10 min at 2095 g and 20°C, 3 mL of the acetonitrile upper layer was collected and introduced in a 15 mL Falcon® tube. The extraction was repeated with 5 mL of acidified acetonitrile and, after centrifugation, a volume of 3 mL acetonitrile was collected and mixed with the previous one. After a night at -20°C (to allow protein precipitation), the 6 mL extracts were centrifuged 10 min at 5250 g and 4°C. The purification step by d-SPE took then place as for soil.

##### 2.5.3 Quantification by liquid chromatography-mass spectrometry

The analysis was performed by on-line Solid Phase Extraction Ultra-High Performance Liquid Chromatography coupled via an ElectroSpray Interface to a triple quadrupole Mass Spectrometer (SPE-UHPLC-ESI-MS/MS), with Acquity Premier 2D and Xevo TQXS equipments (Waters). The SPE preconcentration was performed using on Oasis HLB cartridge (20 x 2.1 mm, granulometry 25 µm, Waters) by injecting 600 µL. The chromatographic separation was obtained with a BEH C18 column (100 x 2.1 mm, granulometry 1.7 mm, Waters, placed in a 30°C oven) and a 0.4 mL/min mobile phase gradient from 100% UP water (A) to 99.9% acetonitrile (B), both solvents being acidified by 0.1% acetic acid. The gradient was: 0 to 1.6 min 100% A (cartridge loading), 1.6 to 20 min change from 100% to 0.1% A (chromatographic analysis), 16 to 20 min 99.9% B (system cleaning) and finally 20 to 24 min back to 100% A and equilibration in starting conditions. In electrospray, source temperature was set at 150°C, desolvation temperature at 550°C, capillary voltage at 3 kV, cone and desolvation gas flow (nitrogen) at 200 and 1000 L/h, respectively. The cone voltage was 6 and 40 V for EPX and IMD, respectively. The detection was performed in positive mode by multiple reaction monitoring, using argon as collision gas (3 x $10^{-3}$ mbar). The transitions used for quantification and confirmation (and collision energy) were respectively m/z 330>101 (36 eV) and m/z 330>121 (18 eV) for EPX, m/z 256>175 (20 eV) and 256>209 (20 eV) for IMD, m/z 334>121 (18 eV) for EPX-d4, m/z 260>179 (18 eV) for IMD-d4 Data acquisition and equipment control were performed using MassLynx software, and data processing was done with QuanLynx (Waters).

To determine pesticide concentrations, standard solutions presenting eight levels of concentration in analytical standards (and fixed concentration in deuterated compounds) were injected at the beginning and end of each series of analyses. It allowed to obtain calibration curves, from the area of the peak of the quantification transition, with a quadratic model and a weighting in 1/X, X being the concentration of the analytical standard. The quality of the analysis was assured by the systematic use of blanks and of control quality samples (checking yields and inter-series variations). The performances of the method in terms of global yields were respectively for EPX and IMD 85% and 40% in soil, and 80% and 67% in earthworms. The limits of quantification of the method (LOQ) were respectively estimated in soil as 0.004 and 0.019 ng/g for EPX and IMD, and in earthworms as 0.2 and 0.8 ng/g.

### 3 Toxicokinetics modeling - Single substances

#### 3.1 Priors used

Priors are the same between models :

- Distrib(kuIMD, LogUniform, 0.0001, 100000); # min, max
- Distrib(keIMD, LogUniform, 0.0001, 100000); # min, max
- Distrib(kperiph, LogUniform, 0.00001, 100000); # min, max
- Distrib(phi, Uniform, 0.0, 1); # min, max
- Distrib(a_growth, TruncNormal, 0.005, 0.0010, -0.5, 1); # mean, sd, min, max
- Distrib (Vr_a_growth, TruncNormal, 0.0015, 0.01, 0, 1); # mean, sd, min, max
- Distrib(Sigma_W, TruncNormal, 0.03, 0.01, 0.0001, 1); # mean, sd, min, max
- Distrib(Sigma_CiIMD, TruncNormal, 3.0, 3.0, 1, 1000); # mean, sd, min, max

Log-Likelihood calculations :

- Likelihood(CiIMD, LogNormal, Prediction(CiIMD), Sigma_CiIMD);
- Likelihood(Weight, Normal, Prediction(Weight), Sigma_W);

#### 3.2 IMD - Model C1

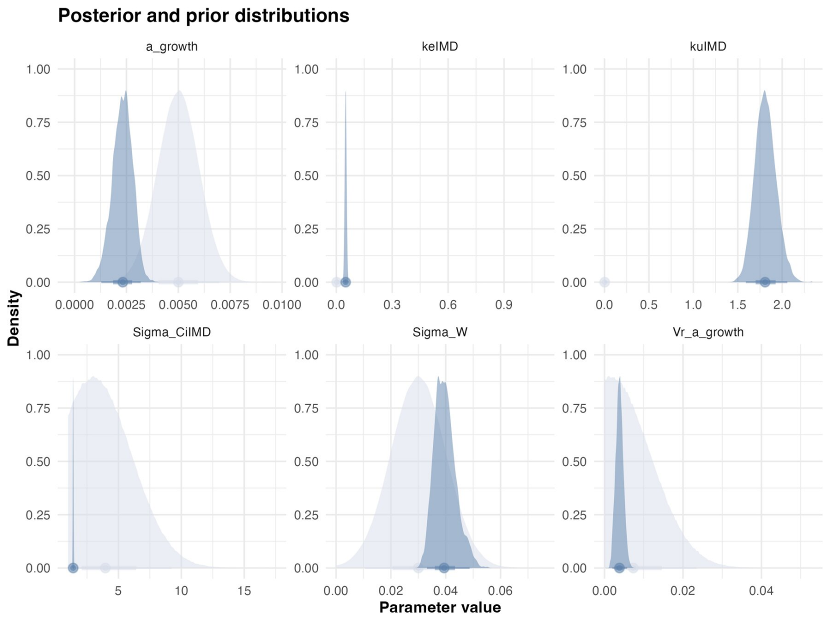

Figure 3.1: **Estimated posterior distributions for each parameter.** Prior and posterior distributions are represented in light and dark blue, respectively.

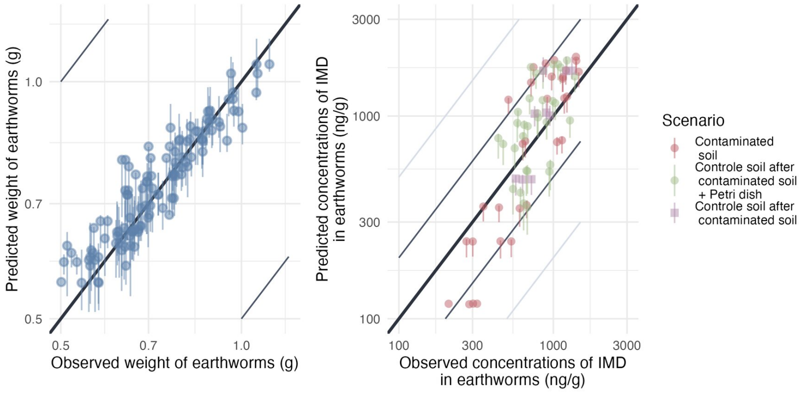

Figure 3.2: **Predicted versus observed values of weight without gut content and internal concentrations.** Points are colored by exposure scenario: red indicates earthworms exposed only to contaminated soil and put on Petri dishes for depuration (uptake phase), green indicates earthworms transferred from contaminated to clean soil and put on Petri dishes for depuration (elimination phase) and violet indicates earthworms exposed only to contaminated soil but only massaged before being frozen. Grey and light-grey lines represent the 2-folds and 5-fold changes, respectively. The bold black line represents the identity line.

Table 3.1: Estimates summary. {tbl-pos='H'}

| Statistic | $k_u$ | $k_e$ | $a_\mu$ | $a_\sigma$ | $W_\sigma$ | $C_{i,\sigma}$ | BAF |
|---|---|---|---|---|---|---|---|
| Mode | 2.0 | 0.050 | 0.0031 | 0.0017 | 0.045 | 1.4 | 36 |
| Q2.5% | 1.6 | 0.041 | 0.0013 | 0.0020 | 0.033 | 1.4 | 32 |
| Q97.5% | 2.1 | 0.060 | 0.0032 | 0.0058 | 0.049 | 1.5 | 41 |

#### 3.3 IMD - Model C1P

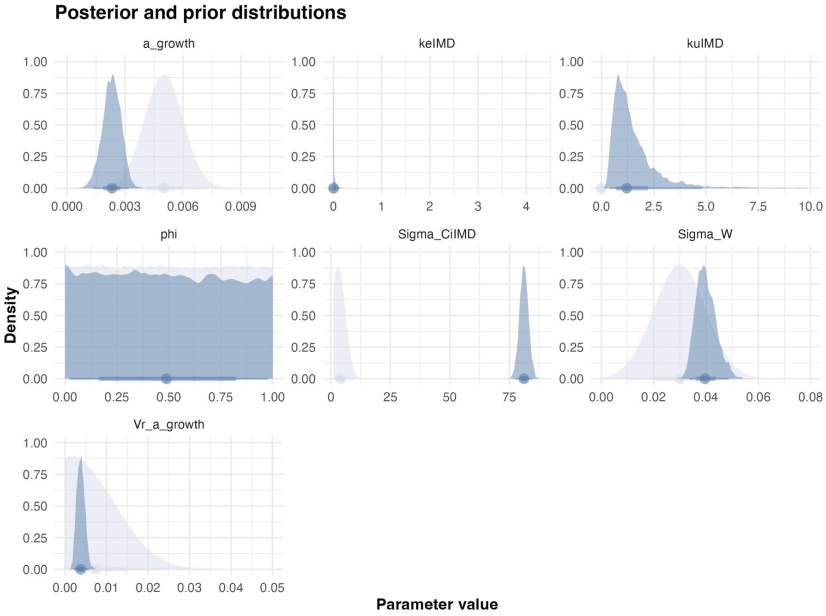

Figure 3.3: **Estimated posterior distributions for each parameter.** Prior and posterior distributions are represented in light and dark blue, respectively.

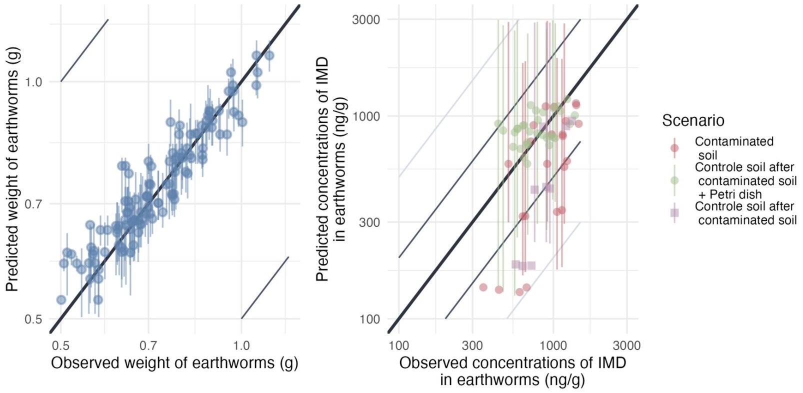

Figure 3.4: **Predicted versus observed values of weight without gut content and internal concentrations.** Points are colored by exposure scenario: red indicates earthworms exposed only to contaminated soil and put on Petri dishes for depuration (uptake phase), green indicates earthworms transferred from contaminated to clean soil and put on Petri dishes for depuration (elimination phase) and violet indicates earthworms exposed only to contaminated soil but only massaged before being frozen. Grey and light-grey lines represent the 2-folds and 5-fold changes, respectively. The bold black line represents the identity line.

Table 3.2: Estimates summary. {tbl-pos='H'}

| Statistic | $k_u$ | $k_e$ | $\phi$ | $a_\mu$ | $a_\sigma$ | $W_\sigma$ | $C_{i,\sigma}$ | BAF |
|---|---|---|---|---|---|---|---|---|
| Mode | 0.68 | 0.00014 | 0.0097 | 0.0023 | 0.0029 | 0.039 | 80 | 3200 |
| Q2.5% | 0.4 | 0.00012 | 0.023 | 0.0013 | 0.0021 | 0.034 | 77 | 17 |
| Q97.5% | 5.1 | 0.12 | 0.97 | 0.0032 | 0.0057 | 0.049 | 85 | 10000 |

#### 3.4 IMD - Model C2N

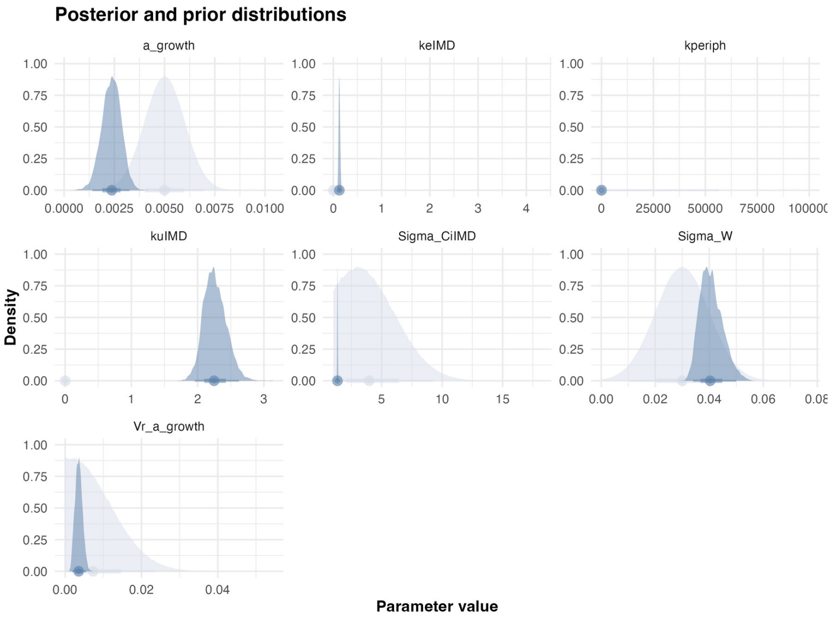

Figure 3.5: **Estimated posterior distributions for each parameter.** Prior and posterior distributions are represented in light and dark blue, respectively.

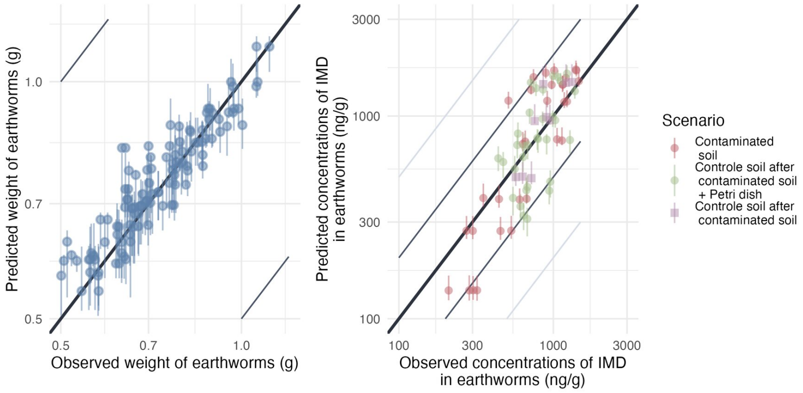

Figure 3.6: **Predicted versus observed values of weight without gut content and internal concentrations.** Points are colored by exposure scenario: red indicates earthworms exposed only to contaminated soil and put on Petri dishes for depuration (uptake phase), green indicates earthworms transferred from contaminated to clean soil and put on Petri dishes for depuration (elimination phase) and violet indicates earthworms exposed only to contaminated soil but only massaged before being frozen. Grey and light-grey lines represent the 2-folds and 5-fold changes, respectively. The bold black line represents the identity line.

Table 3.3: Estimates summary.

| Statistic | $k_u$ | $k_e$ | $k_{periph}$ | $a_\mu$ | $a_\sigma$ | $W_\sigma$ | $C_{i,\sigma}$ | BAF |
|---|---|---|---|---|---|---|---|---|
| Mode | 2.2 | 0.13 | 0.058 | 0.0033 | 0.0018 | 0.046 | 1.3 | 18 |
| Q2.5% | 2.0 | 0.010 | 0.030 | 0.0014 | 0.0018 | 0.034 | 1.3 | 16 |
| Q97.5% | 2.6 | 0.16 | 0.073 | 0.0033 | 0.0055 | 0.050 | 1.4 | 21 |

#### 3.5 IMD - Model C2

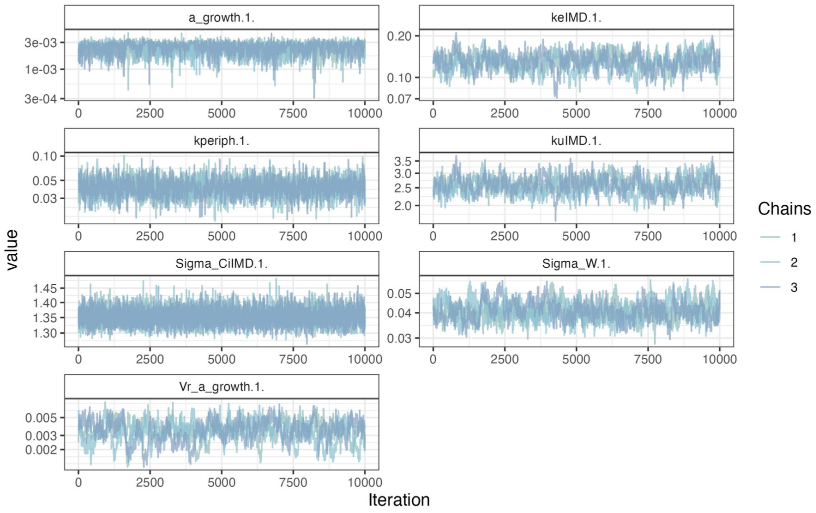

Figure 3.7: **MCMC chains of the model.**

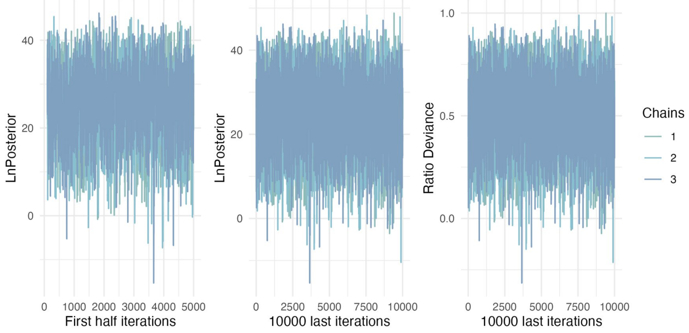

Figure 3.8: **Likelihood of chains.**

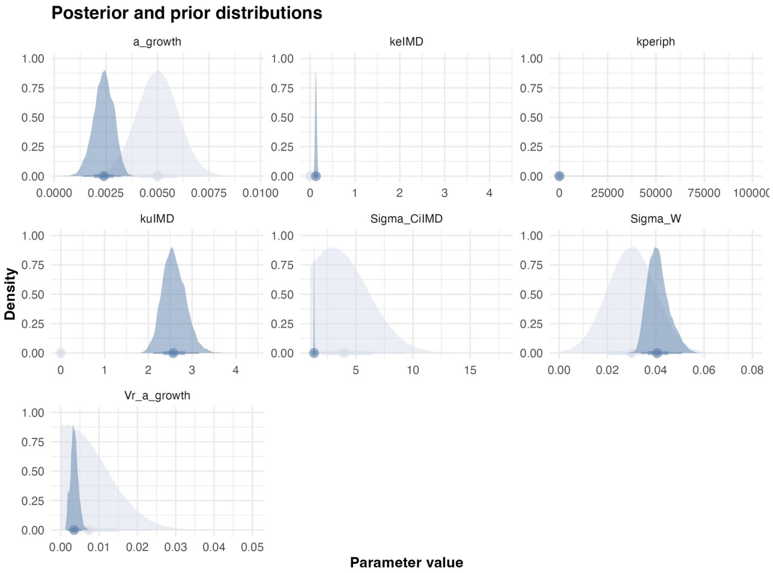

Figure 3.9: **Estimated posterior distributions for each parameter.** Prior and posterior distributions are represented in light and dark blue, respectively.

Table 3.4: Estimates summary.

| Statistic | $k_u$ | $k_e$ | $k_{periph}$ | $a_\mu$ | $a_\sigma$ | $W_\sigma$ | $C_{i,\sigma}$ | BAF |
|---|---|---|---|---|---|---|---|---|
| Mode | 2.4 | 0.12 | 0.045 | 0.0026 | 0.0030 | 0.039 | 1.4 | 21 |
| Q2.5% | 2.1 | 0.096 | 0.025 | 0.0014 | 0.0017 | 0.034 | 1.3 | 18 |
| Q97.5% | 3.2 | 0.17 | 0.063 | 0.0033 | 0.0056 | 0.051 | 1.4 | 23 |

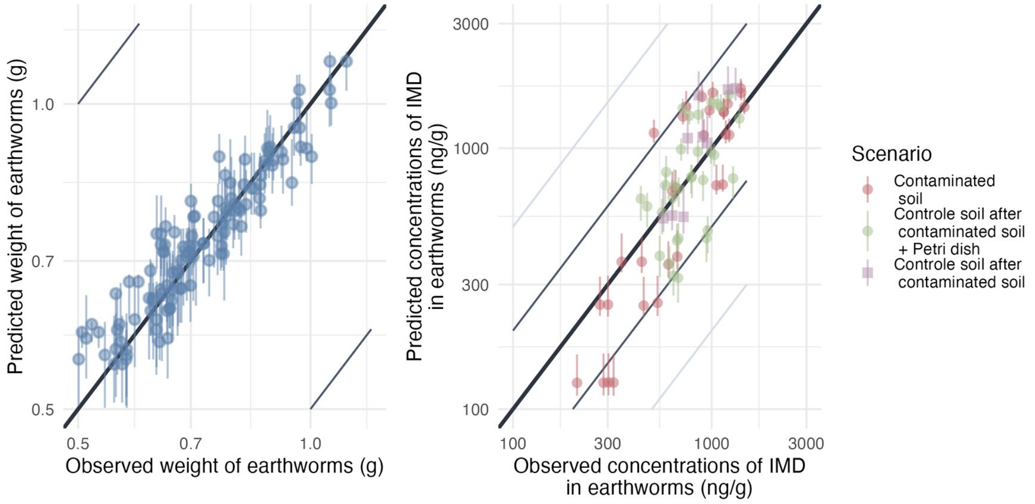

Figure 3.10: **Predicted versus observed values of weight without gut content and internal concentrations.** Points are colored by exposure scenario: red indicates earthworms exposed only to contaminated soil and put on Petri dishes for depuration (uptake phase), green indicates earthworms transferred from contaminated to clean soil and put on Petri dishes for depuration (elimination phase) and violet indicates earthworms exposed only to contaminated soil but only massaged before being frozen. Grey and light-grey lines represent the 2-folds and 5-fold changes, respectively. The bold black line represents the identity line.

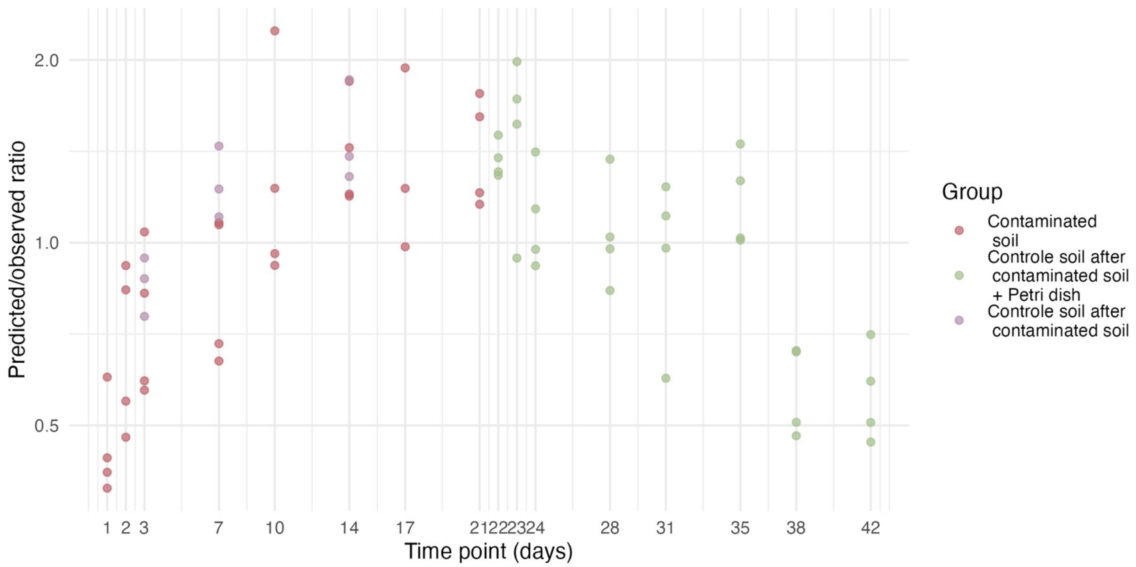

Figure 3.11: **Model predicted/observed ratios.**

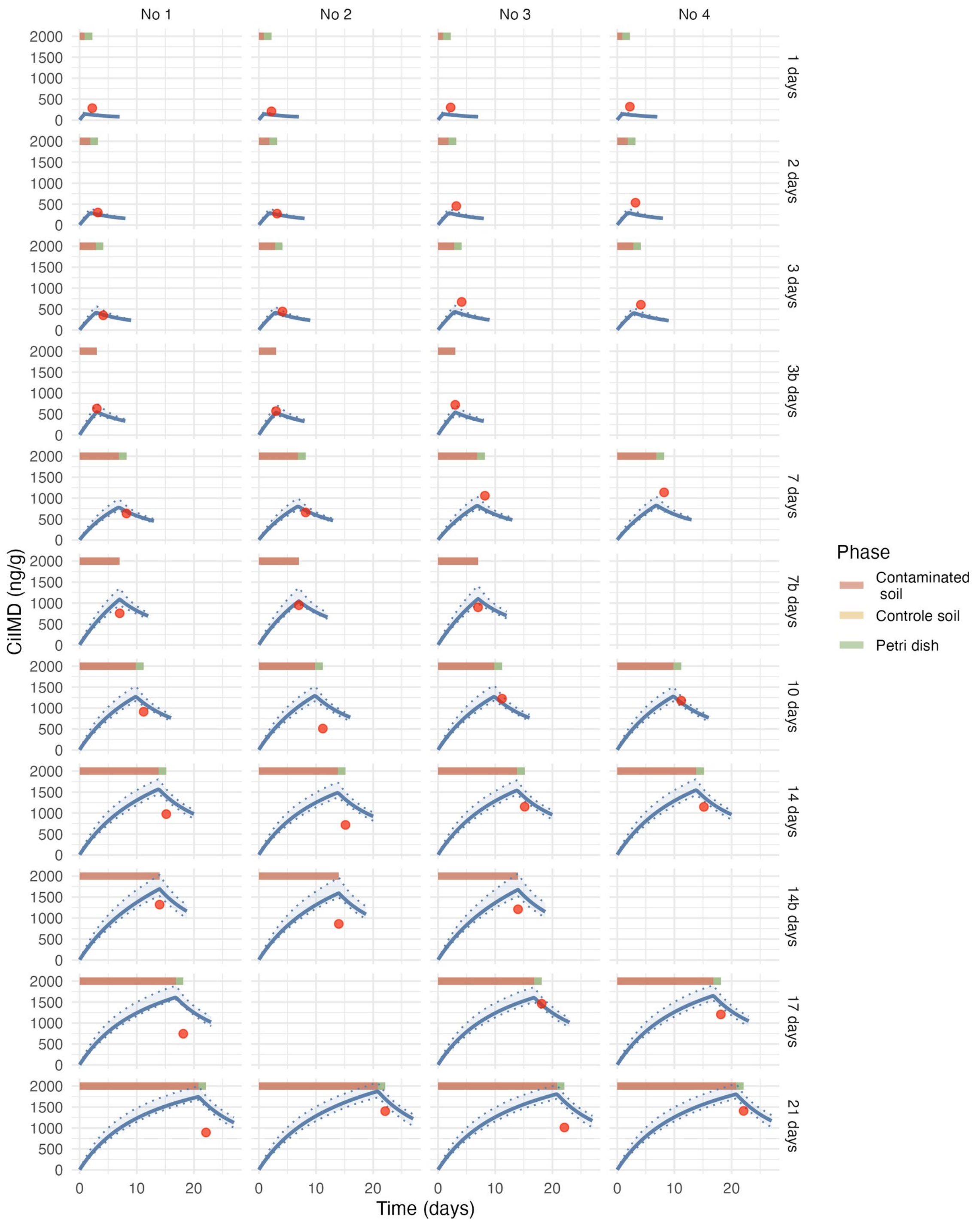

Figure 3.12: **Estimated toxicokinetic of the IMD concentration in earthworms exposed to 1000 ng/g of IMD.** Red points represent the experimental data, the blue curve represent the estimated toxicokinetic curve and the transparent blue band correspond to the credibility interval of the model.

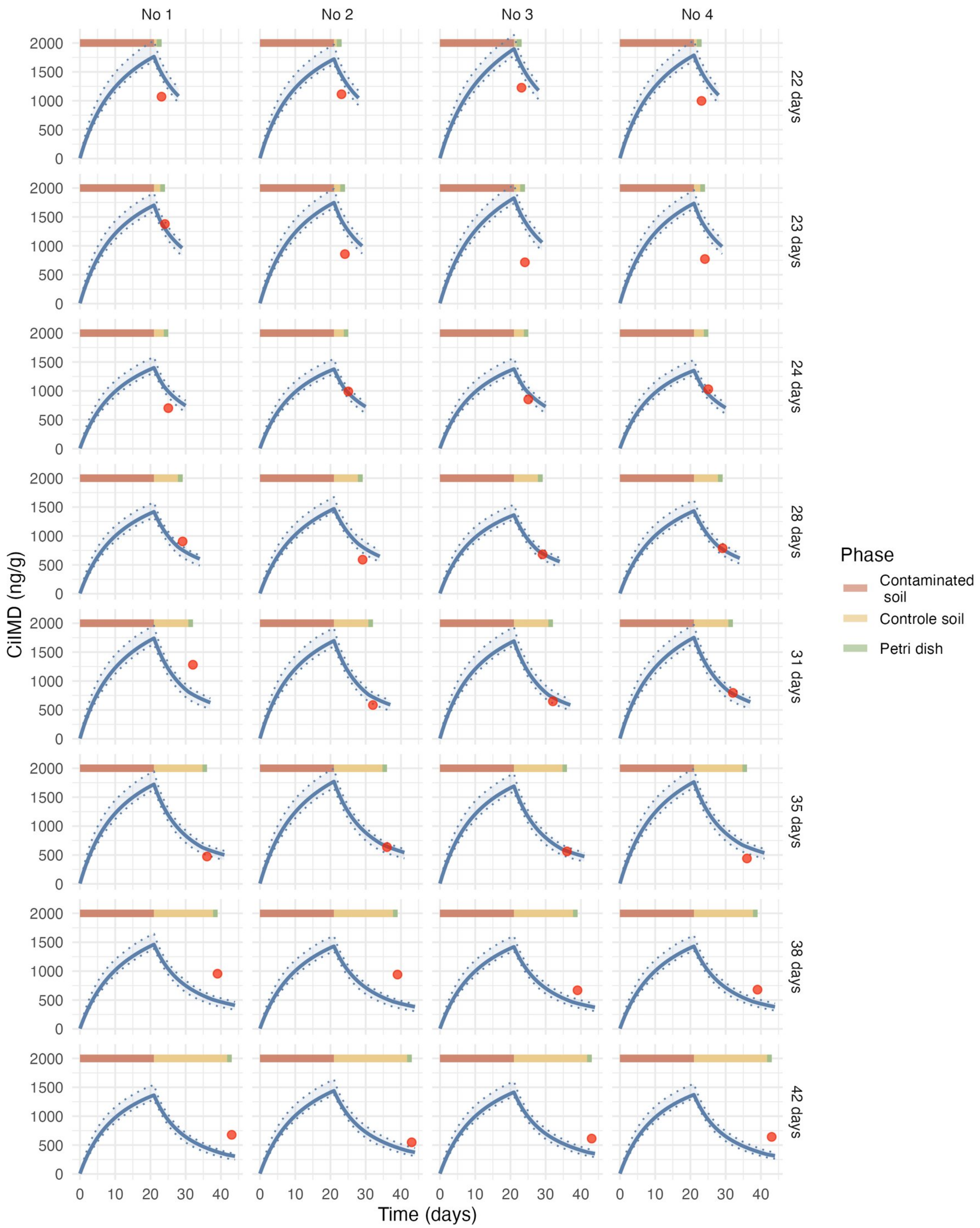

Figure 3.13: **Estimated toxicokinetic of the IMD concentration in earthworms exposed to 1000 ng/g of IMD.** Red points represent the experimental data, the blue curve represent the estimated toxicokinetic curve and the transparent blue band correspond to the credibility interval of the model.

#### 3.6 IMD - Model C2P

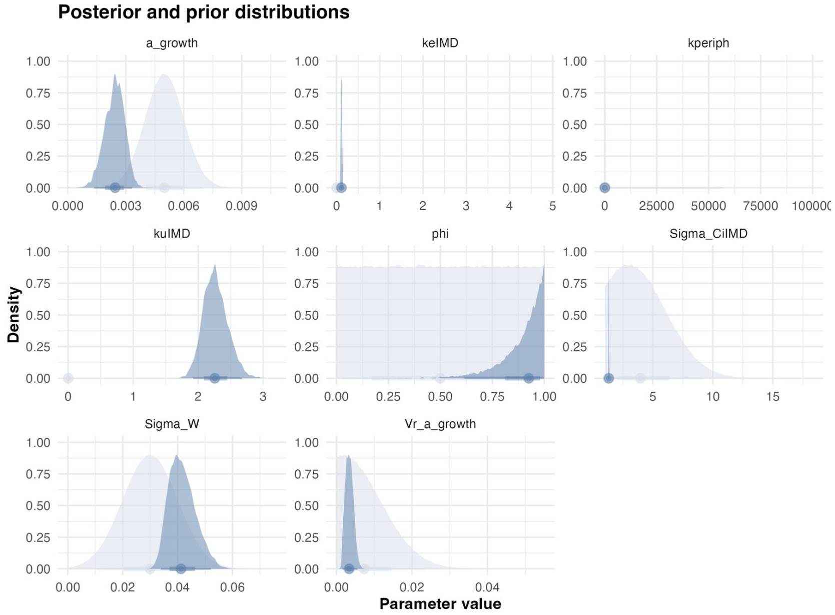

Figure 3.14: **Estimated posterior distributions for each parameter.** Prior and posterior distributions are represented in light and dark blue, respectively.

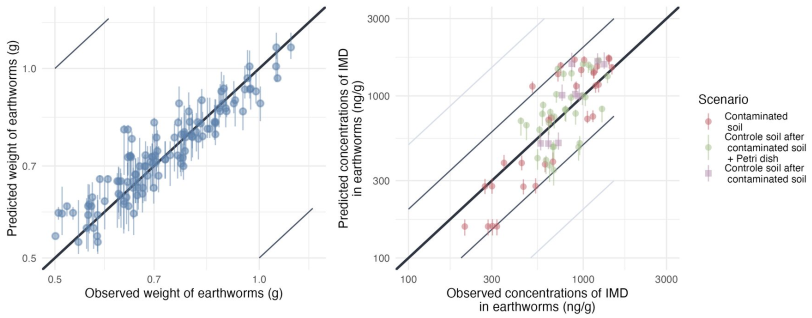

Figure 3.15: **Predicted versus observed values of weight without gut content and internal concentrations.** Points are colored by exposure scenario: red indicates earthworms exposed only to contaminated soil and put on Petri dishes for depuration (uptake phase), green indicates earthworms transferred from contaminated to clean soil and put on Petri dishes for depuration (elimination phase) and violet indicates earthworms exposed only to contaminated soil but only massaged before being frozen. Grey and light-grey lines represent the 2-folds and 5-fold changes, respectively. The bold black line represents the identity line.

Table 3.5: Estimates summary.

| Statistic | $k_u$ | $k_e$ | $k_{periph}$ | $\phi$ | $a_\mu$ | $a_\sigma$ | $W_\sigma$ | $C_{i,\sigma}$ | BAF |
|---|---|---|---|---|---|---|---|---|---|
| Mode | 2.2 | 0.11 | 0.042 | 0.97 | 0.0024 | 0.002 | 0.045 | 1.3 | 22 |
| Q2.5% | 1.9 | 0.087 | 0.025 | 0.62 | 0.0014 | 0.0016 | 0.034 | 1.3 | 18 |
| Q97.5% | 2.7 | 0.14 | 0.065 | 1 | 0.0033 | 0.0056 | 0.052 | 1.4 | 23 |

#### 3.7 EPX - Model C1

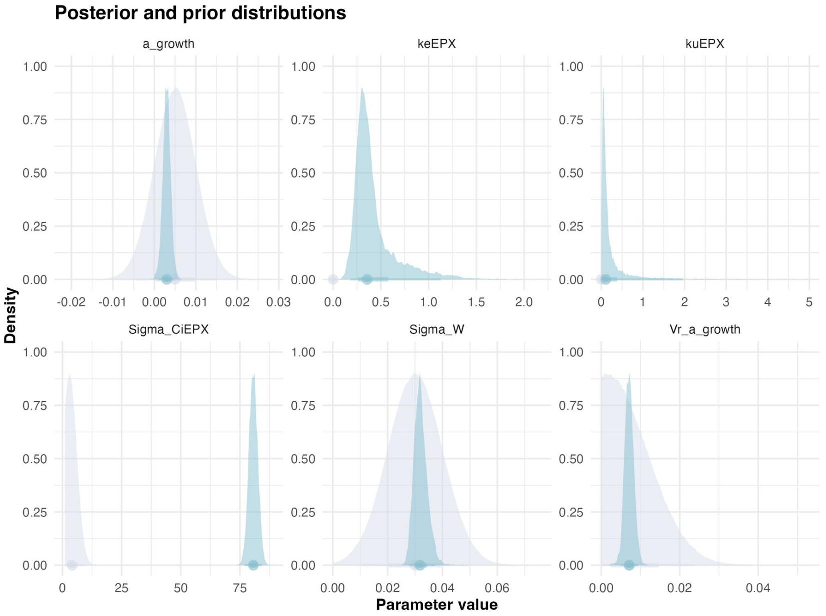

Figure 3.16: **Estimated posterior distributions for each parameter.** Prior and posterior distributions are represented in light and dark blue, respectively.

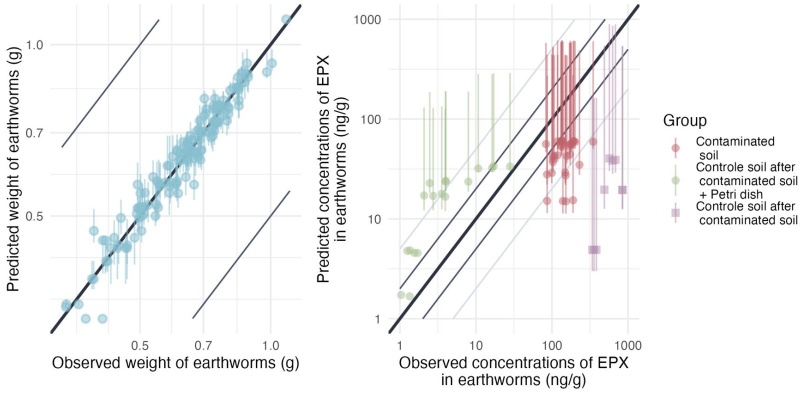

Figure 3.17: **Predicted versus observed values of weight without gut content and internal concentrations.** Points are colored by exposure scenario: red indicates earthworms exposed only to contaminated soil and put on Petri dishes for depuration (uptake phase), green indicates earthworms transferred from contaminated to clean soil and put on Petri dishes for depuration (elimination phase) and violet indicates earthworms exposed only to contaminated soil but only massaged before being frozen. Grey and light-grey lines represent the 2-folds and 5-fold changes, respectively. The bold black line represents the identity line.

Table 3.6: Estimates summary.

| Statistic | $k_u$ | $k_e$ | $a_\mu$ | $a_\sigma$ | $W_\sigma$ | $C_{i,\sigma}$ | BAF |
|---|---|---|---|---|---|---|---|
| Mode | 0.028 | 0.32 | 0.0039 | 0.0076 | 0.028 | 79 | 0.097 |
| Q2.5% | 0.018 | 0.18 | 0.0010 | 0.0049 | 0.028 | 77 | 0.079 |
| Q97.5% | 2.0 | 1.1 | 0.0048 | 0.0095 | 0.037 | 84 | 2 |

#### 3.8 EPX - Model C1P

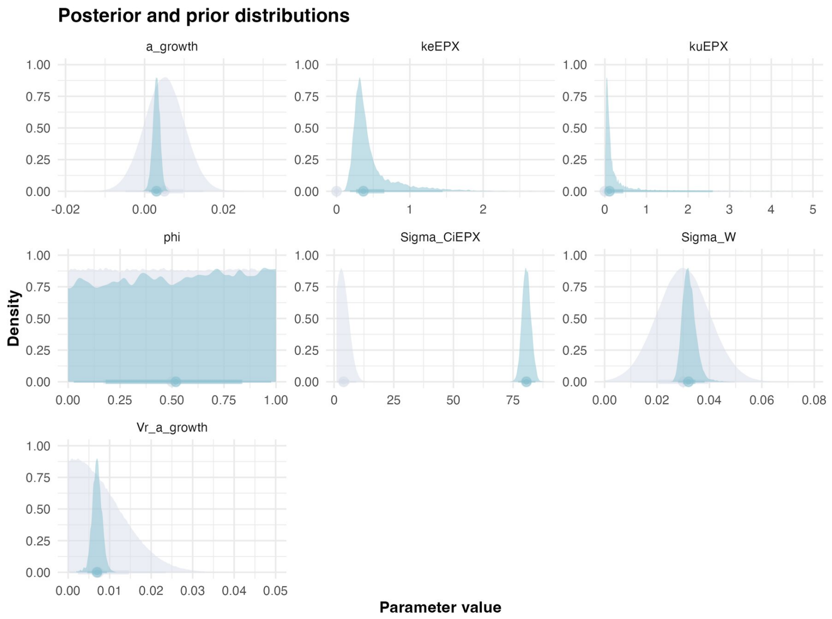

Figure 3.18: **Estimated posterior distributions for each parameter.** Prior and posterior distributions are represented in light and dark blue, respectively.

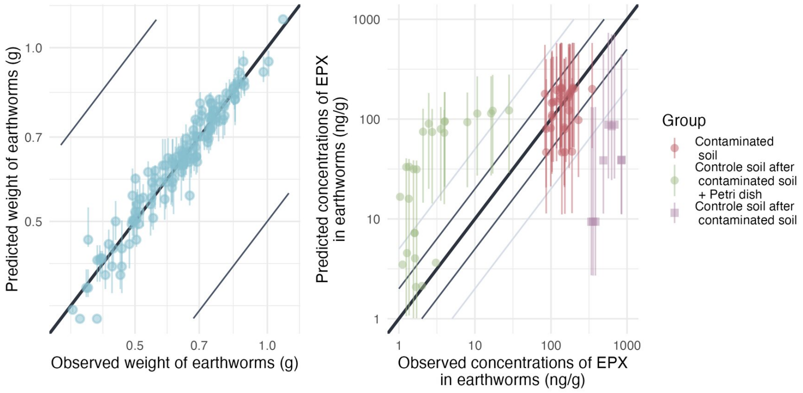

Figure 3.19: **Predicted versus observed values of weight without gut content and internal concentrations.** Points are colored by exposure scenario: red indicates earthworms exposed only to contaminated soil and put on Petri dishes for depuration (uptake phase), green indicates earthworms transferred from contaminated to clean soil and put on Petri dishes for depuration (elimination phase) and violet indicates earthworms exposed only to contaminated soil but only massaged before being frozen. Grey and light-grey lines represent the 2-folds and 5-fold changes, respectively. The bold black line represents the identity line.

Table 3.7: Estimates summary.

| Statistic | $k_u$ | $k_e$ | $\phi$ | $a_\mu$ | $a_\sigma$ | $W_\sigma$ | $C_{i,\sigma}$ | BAF |
|---|---|---|---|---|---|---|---|---|
| Mode | 0.054 | 0.21 | 0.99 | 0.0022 | 0.0076 | 0.030 | 82 | 0.18 |
| Q2.5% | 0.021 | 0.18 | 0.028 | 0.0011 | 0.0046 | 0.028 | 77 | 0.0846 |
| Q97.5% | 3.5 | 1.4 | 0.98 | 0.0047 | 0.0093 | 0.038 | 84 | 2.6 |

#### 3.9 EPX - Model C2

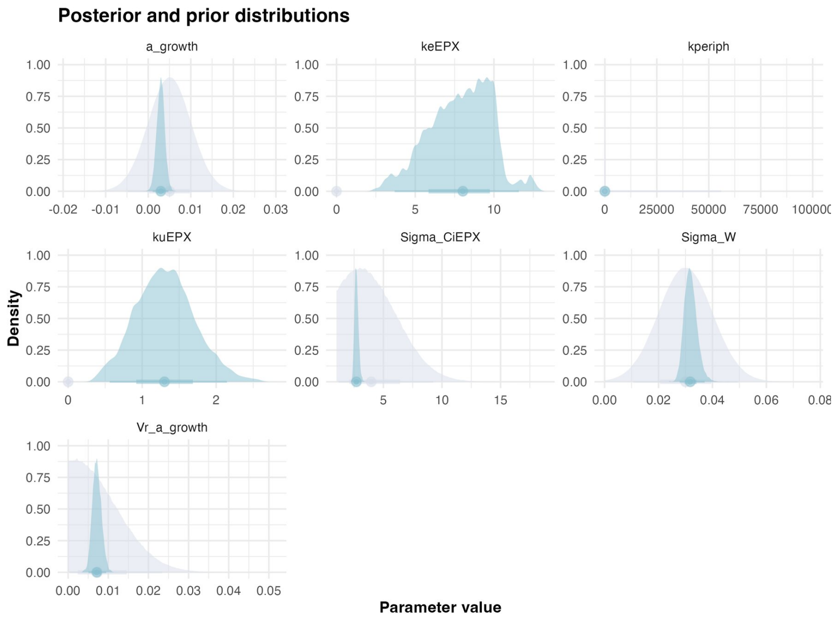

Figure 3.20: **Estimated posterior distributions for each parameter.** Prior and posterior distributions are represented in light and dark blue, respectively.

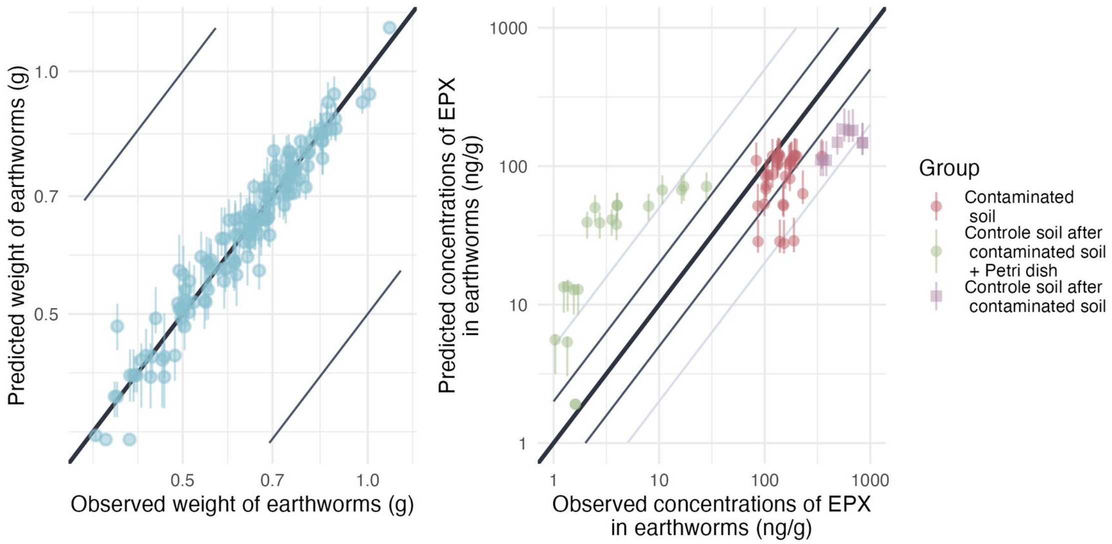

{fig-scap="Predicted versus observed values of internal concentrations. fig-pos="H" width="70%"}

Table 3.8: Estimates summary.

| Statistic | $k_u$ | $k_e$ | $k_{periph}$ | $a_\mu$ | $a_\sigma$ | $W_\sigma$ | $C_{i,\sigma}$ | BAF |
|---|---|---|---|---|---|---|---|---|
| Mode | 1.3 | 8.3 | 0.28 | 0.0021 | 0.0083 | 0.029 | 2.5 | 0.16 |
| Q2.5% | 0.56 | 3.7 | 0.27 | 0.0010 | 0.0051 | 0.028 | 2.4 | 0.12 |
| Q97.5% | 2.1 | 12.0 | 0.35 | 0.0048 | 0.0095 | 0.037 | 3.1 | 0.22 |

#### 3.10 EPX - Model C2P

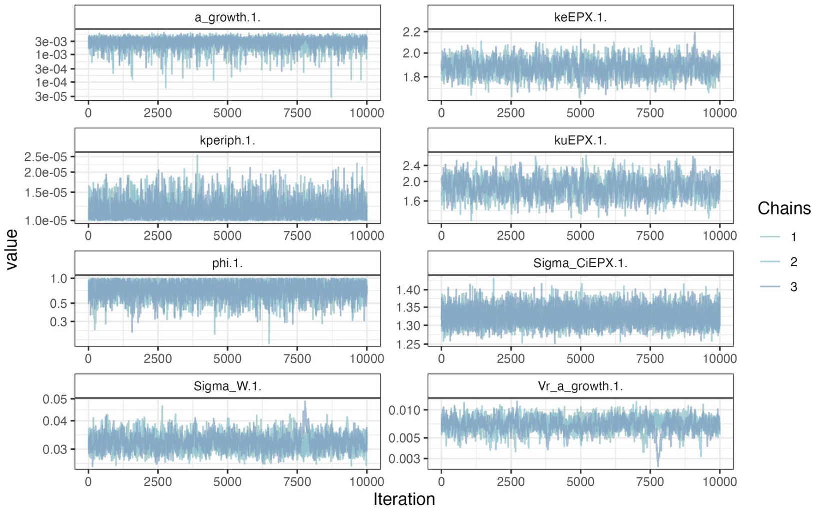

Figure 3.21: **MCMC chains of the model.**

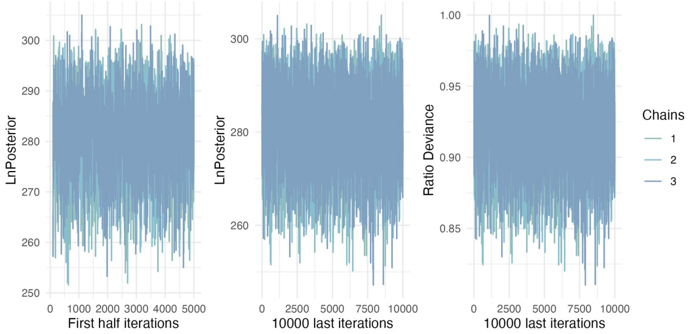

Figure 3.22: **Likelihood of chains.**

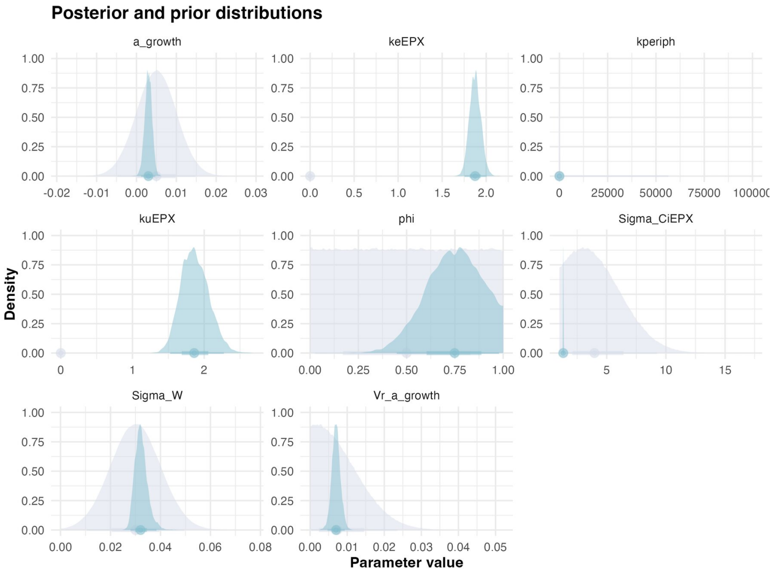

Figure 3.23: **Estimated posterior distributions for each parameter.** Prior and posterior distributions are represented in light and dark blue, respectively.

Table 3.9: Estimates summary.

| Statistic | $k_u$ | $k_e$ | $k_{periph}$ | $\phi$ | $a_\mu$ | $a_\sigma$ | $W_\sigma$ | $C_{i,\sigma}$ | BAF |
|---|---|---|---|---|---|---|---|---|---|
| Mode | 1.8 | 1.8 | 0.000010 | 0.67 | 0.0037 | 0.0076 | 0.028 | 1.3 | 0.87 |
| Q2.5% | 1.5 | 1.8 | 0.000010 | 0.45 | 0.0011 | 0.0047 | 0.028 | 1.3 | 0.85 |
| Q97.5% | 2.3 | 2.10 | 0.000015 | 0.98 | 0.0047 | 0.0093 | 0.038 | 1.4 | 1.2 |

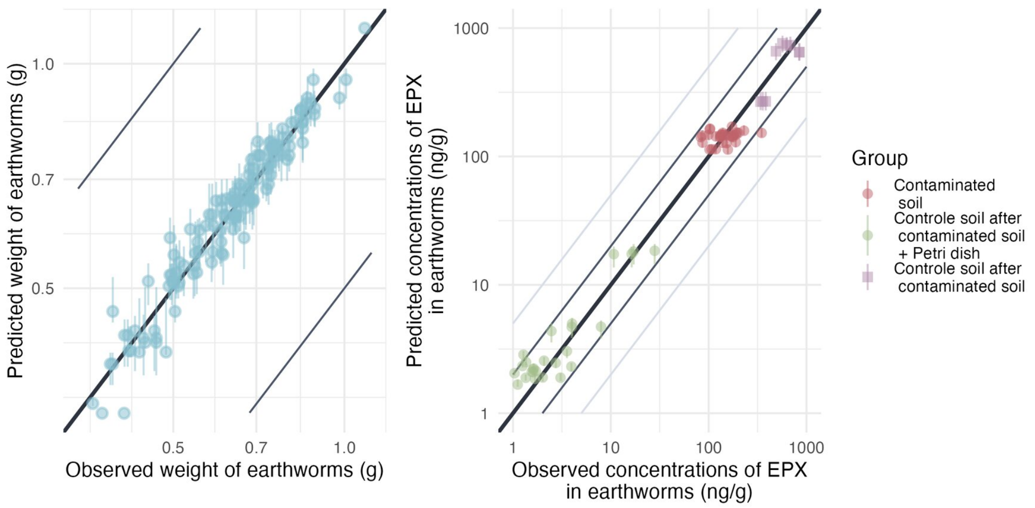

Figure 3.24: **Predicted versus observed values of weight without gut content and internal concentrations.** Points are colored by exposure scenario: red indicates earthworms exposed only to contaminated soil and put on Petri dishes for depuration (uptake phase), green indicates earthworms transferred from contaminated to clean soil and put on Petri dishes for depuration (elimination phase) and violet indicates earthworms exposed only to contaminated soil but only massaged before being frozen. Grey and light-grey lines represent the 2-folds and 5-fold changes, respectively. The bold black line represents the identity line.

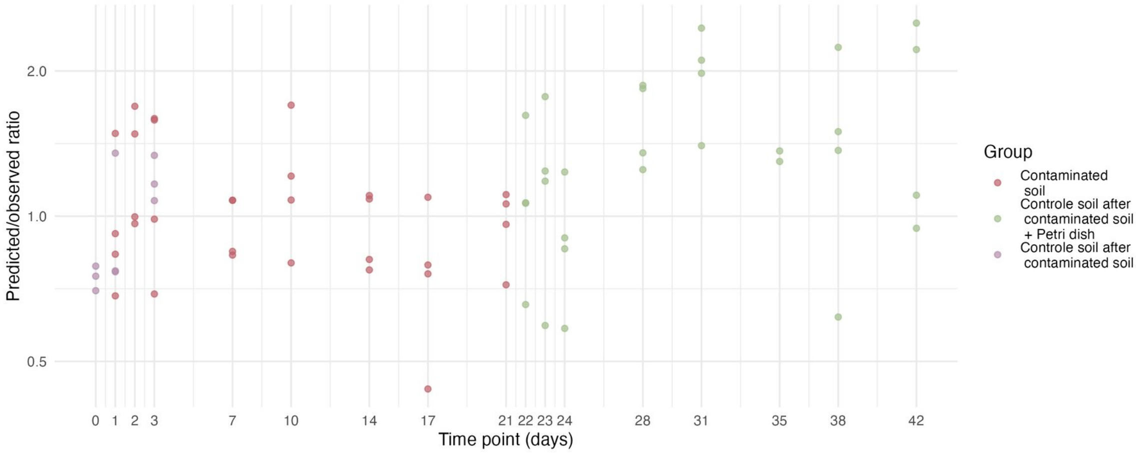

Figure 3.25: Model predicted/observed ratios.

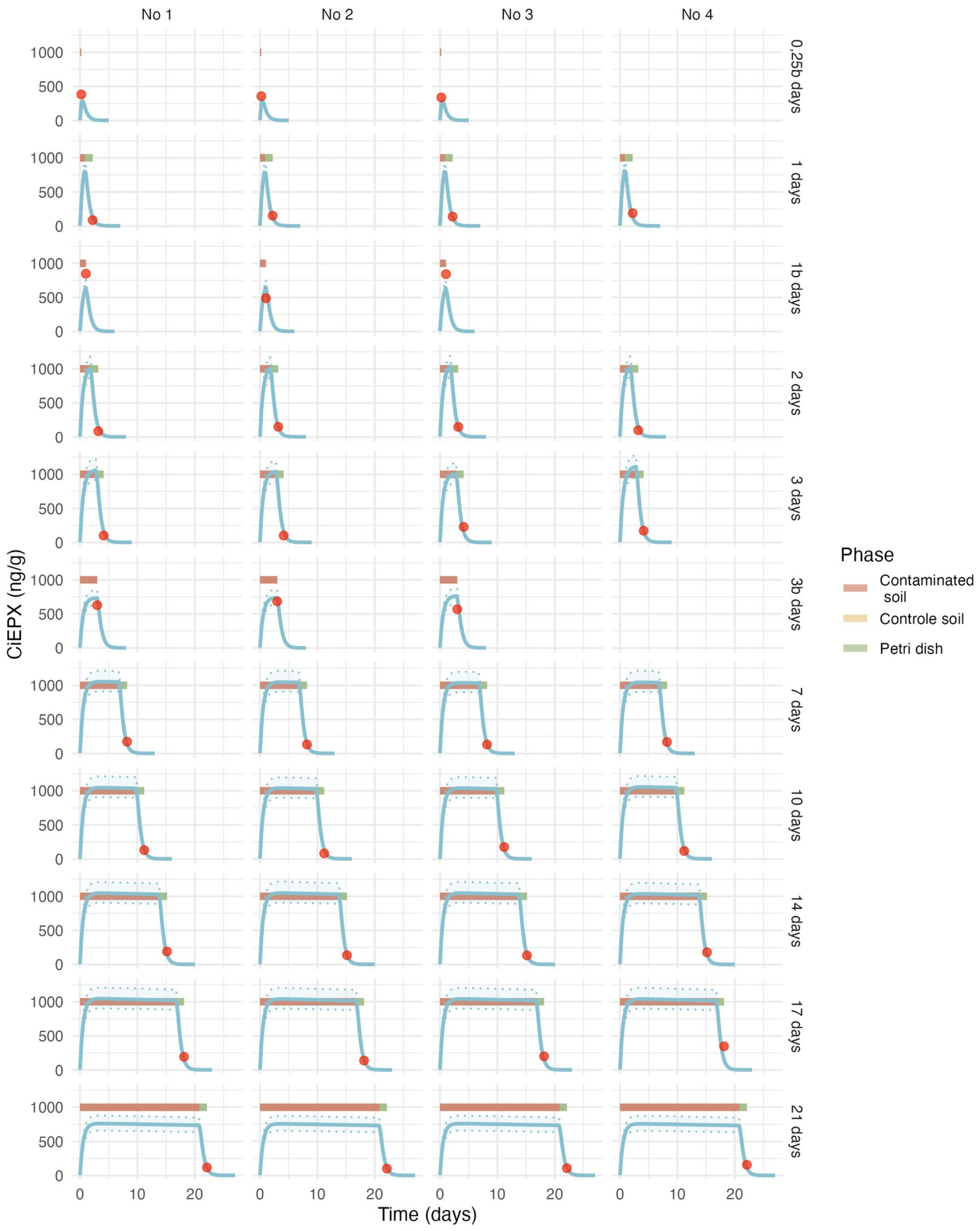

Figure 3.26: Estimated toxicokinetic of the EPX concentration in earthworms exposed to 1000 ng/g of EPX. Red points represent the experimental data, the blue curve represent the estimated toxicokinetic curve and the transparent blue band correspond to the credibility interval of the model.

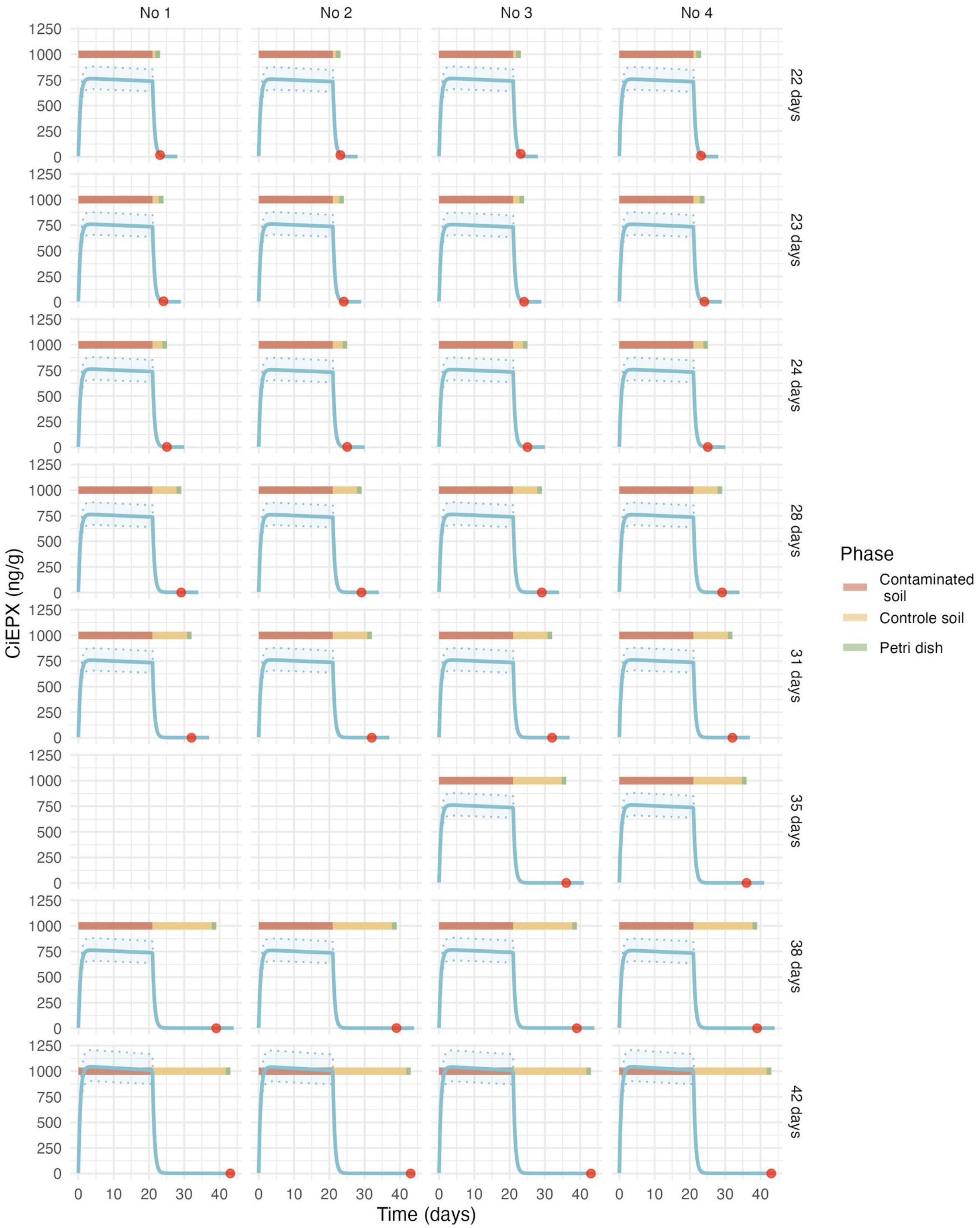

Figure 3.27: Estimated toxicokinetic of the EPX concentration in earthworms exposed to 1000 ng/g of EPX. Red points represent the experimental data, the blue curve represent the estimated toxicokinetic curve and the transparent blue band correspond to the credibility interval of the model.

Bart, S., Laurent, C., Péry, A. R. R., Mougin, C., & Pelosi, C. (2017). Differences in sensitivity between earthworms and enchytraeids exposed to two commercial fungicides. *Ecotoxicology and Environmental Safety*, *140*, 177–184. https://doi.org/10.1016/j.ecoenv.2017.02.052

Kabus, J., Hartmann, V., Cocchiararo, B., Dombrowski, A., Enns, D., Karaouzas, I., Lipkowski, K., Pelikan, L., Shumka, S., Soose, L., Baker, N. J., & Jourdan, J. (2024). Cryptic species complex shows population-dependent, rather than lineage-dependent tolerance to a neonicotinoid. *Environmental Pollution*, *362*, 124888. https://doi.org/10.1016/j.envpol.2024.124888
